# Th1 effector CD4 T cells rely on IFN-γ production to induce alopecia areata

**DOI:** 10.1101/2025.05.07.650678

**Authors:** Samuel J. Connell, Sydney Crotts, Ryan Reis, Maddison Lensing, Payton Kahl, Nicholas Henderson, Otgonzaya Ayush, Zhaowen Zhu, Luana S. Ortolan, John Harty, Joan Goverman, Ali Jabbari

**Author notes:** Corresponding Author: Department of Dermatology, University of Iowa, 2110 Medical Laboratories, 25 S. Grand Ave, Iowa City, IA, 52242.

## Abstract

Alopecia areata (AA) is an autoimmune disease that is clinically characterized by hair loss and histologically by a peribulbar infiltrate of CD8 and CD4 T cells. Prior studies have focused on the role of CD8 T cells in the development of AA; however, the role of CD4 T cells remains unclear. Here, we demonstrate that CD4 T cells from the skin draining lymph nodes (SDLN) of AA mice transferred disease into recipient mice. Further, these cells exhibited a T-helper type 1 (Th1) effector transcriptional and phenotypic profile. The pathogenic activity of these CD4 T cells was dependent upon the presence of endogenous CD8 T cells and host IFN-γ responsiveness. Targeted deletion of CD4 T cell-mediated production of IFN-γ abrogated the ability of this cell population to transfer disease. Together, these data provide mechanistic insights into pathways that lead to AA development, strengthening our understanding of the disease and inviting studies into exploring novel therapeutic strategies for human patients.

## Introduction

Alopecia areata (AA) is an autoimmune disease of the hair follicle with a lifetime incidence rate of approximately 2% (Mirzoyev et al., 2014, Safavi et al., 1995, Mostaghimi et al., 2023). Clinical presentation of AA can range from small, localized patches of hair loss to widespread hair loss of the whole body (Finner, 2011, Gilhar et al., 2007). Recently, the FDA approved the use of Janus Kinase (JAK) inhibitors for the treatment of AA. However, this family of drugs have shown incomplete efficacy in clinical trials, and use of JAK inhibitors has the potential for severe side effects including opportunistic infections, cancer, and, thrombosis (Hoisnard et al., 2022, Ytterberg et al., 2022, King et al., 2022, King et al., 2024, King et al., 2023). Further, following discontinuation of treatment, many patients often undergo a relapse of disease and lose hair. These caveats support the need to continue the investigation into the immune mechanisms involved in AA.

The hair follicle is considered an immune privileged (IP) site. Contributing factors to this state include minimal expression of major histocompatibility complex (MHC) molecules in the lower parts of the follicle, a lack of immune cell infiltration, and the presence and action of immunoregulatory factors (Bertolini et al., 2020). AA is thought to occur due to the collapse of IP of the hair follicle, indicated by an upregulation of the expression of MHC molecules and danger signals involving the epithelium of the hair follicle. This loss of IP is thought to be followed by infiltration of autoreactive immune cells into the hair follicle environment, which then perform effector functions that target the lower regions of the follicle and result in non-scarring hair loss (Bertolini et al., 2020, Connell and Jabbari, 2022, Lensing and Jabbari, 2022). The immune cell infiltrate is largely comprised of T cells, including both CD4 and CD8 T cells. Prior work has highlighted a pivotal role for cytotoxic CD8 T cells that express NKG2D in AA pathogenesis (Xing et al., 2014, Petukhova et al., 2010). Examination of lesional scalp tissues from patients with AA have observed that CD4 T cells outnumber CD8 T cells in the T cell infiltrate (Todes-Taylor et al., 1984, Ito et al., 2008, Ito et al., 2013); however, their contributions to the development of AA is not well understood.

Prior work using the C3H/HeJ AA mouse model demonstrated that transfer of cells derived from the skin draining lymph node (SDLN) of AA mice is sufficient to induce disease in recipient mice (McElwee et al., 2005, Wang et al., 2015); subsequent studies supported that sorted CD8 T cells are able to transfer disease (Xing et al., 2014, Dai et al., 2021, Seok et al., 2023). CD4 T cells from AA donor mice have been shown to induce disease in recipient mice when transferred subcutaneously (McElwee et al., 2005). However, the specific contributions of CD4 T cells in the pathogenesis of AA has not yet been defined. Given their presence in AA lesional patient samples and their potential to induce disease in the murine model, our aim was to identify the mechanisms used, and factors required, for CD4 T cell-mediated effects in AA.

Here, we demonstrate that CD4 T cells from AA mice have a robust ability to induce disease when adoptively transferred into recipient mice. Focused examination of CD4 T cells in AA SDLNs revealed that this population exhibited a heightened activated, T-helper type 1 (Th1)- effector gene profile when compared to CD4 T cells from unaffected mice. Additionally, CD4 T cell-mediated disease induction was dependent upon the presence of endogenous CD8 T cells and host interferon gamma (IFN-γ) signaling. Single-cell RNA sequencing (scRNA-seq) data of human scalp tissue revealed that CD4 T cells in AA patients had an analogous, Th1 gene-enriched profile. Disruption of CD4 T cell-mediated production of IFN-γ abolished the ability of CD4 T cells to transfer disease. In sum, we found that Th1-associated effector T cells serve a pivotal role in the development of AA.

## Results

### SDLN from AA mice contain CD8 and CD4 T cells that can contribute to disease induction

It has been previously demonstrated that lymphocytes from the SDLN of mice with AA induce disease in recipient mice (McElwee et al., 2005, Wang et al., 2015). We adopted a model in which lymphocytes from AA SDLNs that underwent *in vitro* expansion (Wang et al., 2015) induced disease when adoptively transferred into recipient mice (Figure 1A). We were interested in defining the composition of the transferred population to gain insights into which specific components may be needed to induce disease. Flow cytometric analysis showed that the cultured cell population comprised of both CD4^+^ and CD8^+^ T cells (Figure 1B), supporting a potential role for either cell type in the induction of AA. An examination of human biopsy specimens demonstrated a robust population of T cells around the hair follicle in AA patients that was not observed in healthy control samples (Figure 1C). These peribulbar T cell populations in AA skin were found to be comprised of predominantly CD4^+^ T cells as well as a smaller population of CD8^+^ T cells (Figure 1D), similar to what others have observed (Ito et al., 2008, Ito et al., 2013). We then performed a transcriptomic analysis of skin samples from healthy control and AA patients. We generated a robust scRNA-seq dataset by compiling data from multiple studies (Borcherding et al., 2020, Lee et al., 2023, Ober-Reynolds et al., 2023), integrated in a manner to mitigate batch effects using established protocols. The dataset was comprised of 11 AA patient samples and 9 healthy control samples consisting of 86,110 total cells, composed of 48,197 from AA patients and 37,913 from control patients. Uniform manifold approximation and projection (UMAP) dimensionality reduction resulted in the identification of clusters representing various immune cell populations (CD4 T cells, CD8 T cells, NK/γδ T cells, myeloid cells, dendritic cells, B cells, and mast cells), skin cell populations (keratinocytes, fibroblasts, and melanocytes), and other various cell types (smooth muscle cells, vascular endothelial cells, lymphatic endothelial cells, and mitotic) (Figure 1E, Supplemental Figure 1).

**Figure 1.**
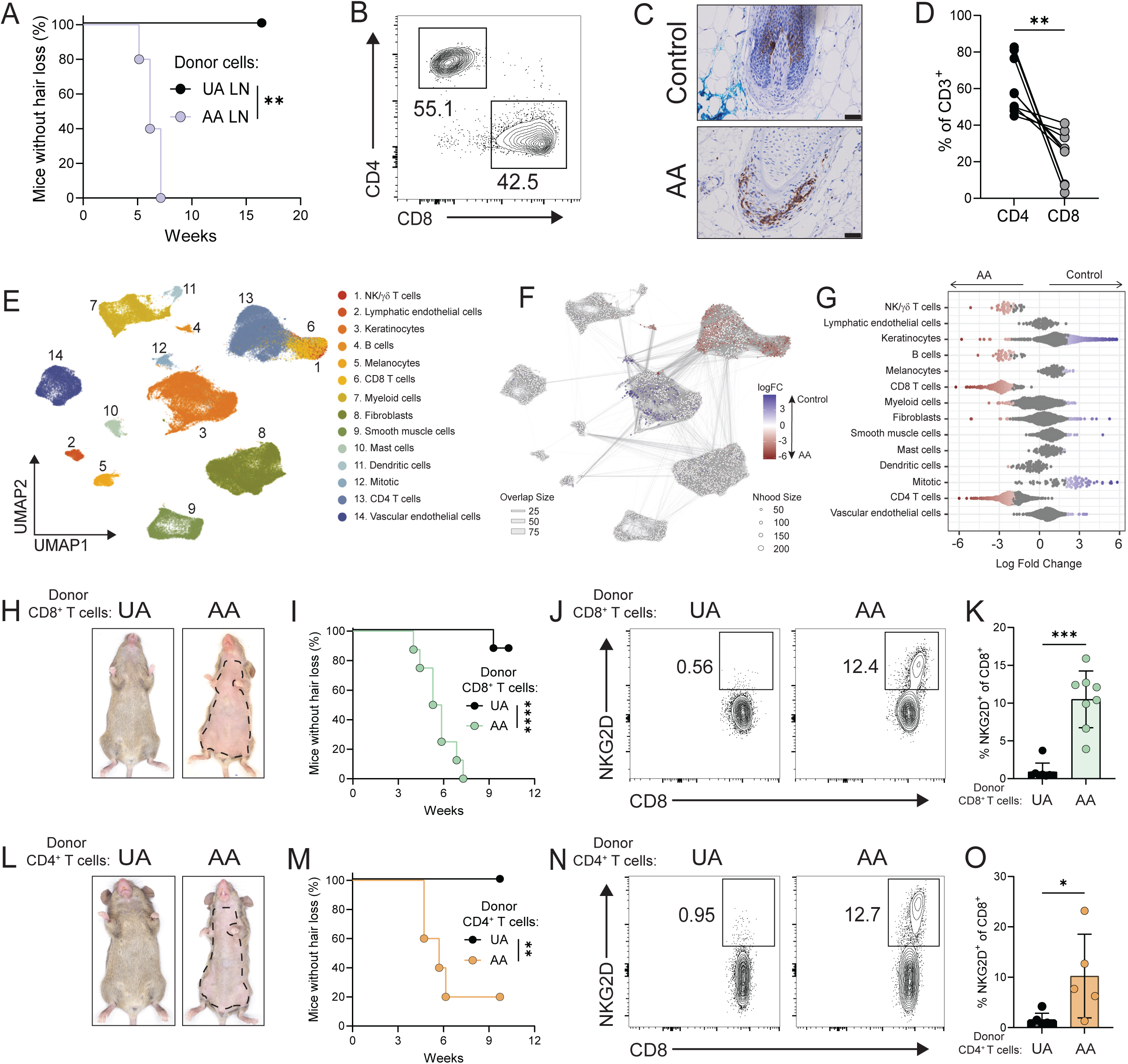
CD8 and CD4 T cells from skin draining lymph nodes (SDLN) of AA mice are sufficient to transfer disease. **(A)** AA development in mice following transfer of *in vitro* expanded bulk SDLN cells from AA and unaffected (UA) donors (n=5 per group). **p=0.0026, Log-rank test. **(B)** Representative flow cytometric analysis of CD4 and CD8 expression from SDLN cells of AA mice following 6 days of *in vitro* expansion. Cells gated on live, CD3^+^ cells. **(C)** Representative immunohistochemical staining of human scalp tissue from healthy control and AA samples showing CD3 protein expression. Scale bar: 100μm. **(D)** Proportion of CD4 and CD8 T cells in CD3^+^ cells from AA human scalp tissue, as assessed by flow cytometry. **p=0.0039, non-parametric paired t-test. **(E)** UMAP of compiled human AA and healthy control scRNA-seq dataset, showing clustering of 86,110 total cells (48,197 AA; 37,913 control). **(F)** Neighborhood graph of the results from Milo differential abundance testing. Nodes are neighborhoods, colored by their log fold change across disease. Non-differential abundance neighborhoods (false discovery rate 10%) are colored white, and sizes correspond to the number of cells in each neighborhood. Lines depict the number of cells shared between neighborhoods. The layout of nodes is determined by the position of the neighborhood index cell in the UMAP in panel E. **(G)** Beeswarm plot of the log-transformed fold-changes in abundance of cells in AA versus control skin. Differential abundance neighborhoods at a false discovery rate of 10% are colored. **(H to K)** Isolated CD8 T cells from SDLNs of UA and AA donor mice underwent *in vitro* expansion and were transferred intradermally into recipient mice (n=8 both groups). **(H & I)** Representative ventral images of mice after receiving intradermal injections and Kaplan-Meier disease curve. ****p<0.0001, Log-Ranked Test. **(J & K)** Representative flow cytometric plots and summary graphs of NKG2D^+^CD8^+^ T cells in the SDLNs of mice (mean ± SD). ***p=0.0002, Mann-Whitney test. **(L to O)** Isolated CD4 T cells from SDLNs of UA and AA donor mice underwent *in vitro* expansion and were transferred intradermally into recipient mice (n=6 UA, n=5 AA). **(L & M)** Representative images of ventral surface of mice after receiving intradermal injections and Kaplan-Meier disease curve. **p=0.0067, Log-rank test. **(N & O)** Representative flow cytometric plots and summary graphs of NKG2D^+^CD8^+^ T cells in the SDLNs of mice (mean ± SD). *p=0.0173, Mann-Whitney test. Mice were observed two times weekly for the development of hair loss. Data is representative of two to three independent experiments.

We performed differential abundance testing using MiloR to interrogate differences in cellular abundance for the broad cell types seen in AA versus control skin (Dann et al., 2022). MiloR was applied to our harmonized dataset, and, using a 10% false discovery rate, we identified 5,914 neighborhoods, with 942 showing an increased abundance in AA skin (shown in red) (Figure 1F). We observed significant enrichment of multiple cell types in AA samples including both CD8 T cells and CD4 T cells (Figure 1G). Together, these data support that both CD8 and CD4 T cells participate in the immune infiltration of lesional AA skin, suggesting that both T cell populations could have a role in the development of disease.

We next investigated the potential of CD8 and CD4 T cells to contribute to the induction of disease in the mouse model. CD8 T cells were magnetically sorted from the SDLNs of unaffected (UA) and AA donor mice. After *in vitro* expansion and transfer to recipient mice, we found that CD8 T cells derived from AA mice induced disease, whereas CD8 T cells from previously unaffected mice were largely unable (Figure 1, H & I). This is consistent with data from other groups that have shown transfer of CD8 T cells from the SDLNs of AA mice, but not from UA mice, can induce disease (McElwee et al., 2005, Xing et al., 2014, Seok et al., 2023). Flow cytometric analysis of the SDLNs of these mice demonstrate that the CD8 T cell-mediated induction is associated with the emergence of NKG2D^+^CD8^+^ T cells (Figure 1, J & K). We next focused on the inductive ability of CD4 T cells. Interestingly, CD4 T cells derived from AA mice had a robust capacity to transfer disease, whereas CD4 T cells derived from UA mice did not (Figure 1, L & M). Additionally, recipients that developed disease following the CD4 T cell transfer demonstrated the development of NKG2D^+^CD8^+^ T cells in the SDLNs (Figure 1, N & O). Disease induction occurred at a faster rate when a larger number of CD4 T cells were transferred (Supplemental Figure 2, A & B). Together, these results demonstrate that CD4 or CD8 T cells from AA mice each have the ability to induce AA.

### AA inductive capacity is restricted to CD4 T cells from the SDLNs

Lymphadenopathy and splenomegaly are features seen in C3H/HeJ mice with AA, but it is unclear which populations are expanded in the setting of AA in the SDLNs and spleen (Xing et al., 2014). We found that mice with AA indeed had an increased number of cells in the spleen (Figure 2A). Further analysis demonstrated there was an increase in the number of multiple immune cell populations including CD11c^+^ cells, αβ T cells, and B cells (Supplemental Figure 3A). Interestingly, within the αβ T cell compartment, we found an increase in the number of CD4 T cells, including both conventional CD4 T cells and regulatory CD4 T cells (Supplemental Figure 3B).

**Figure 2.**
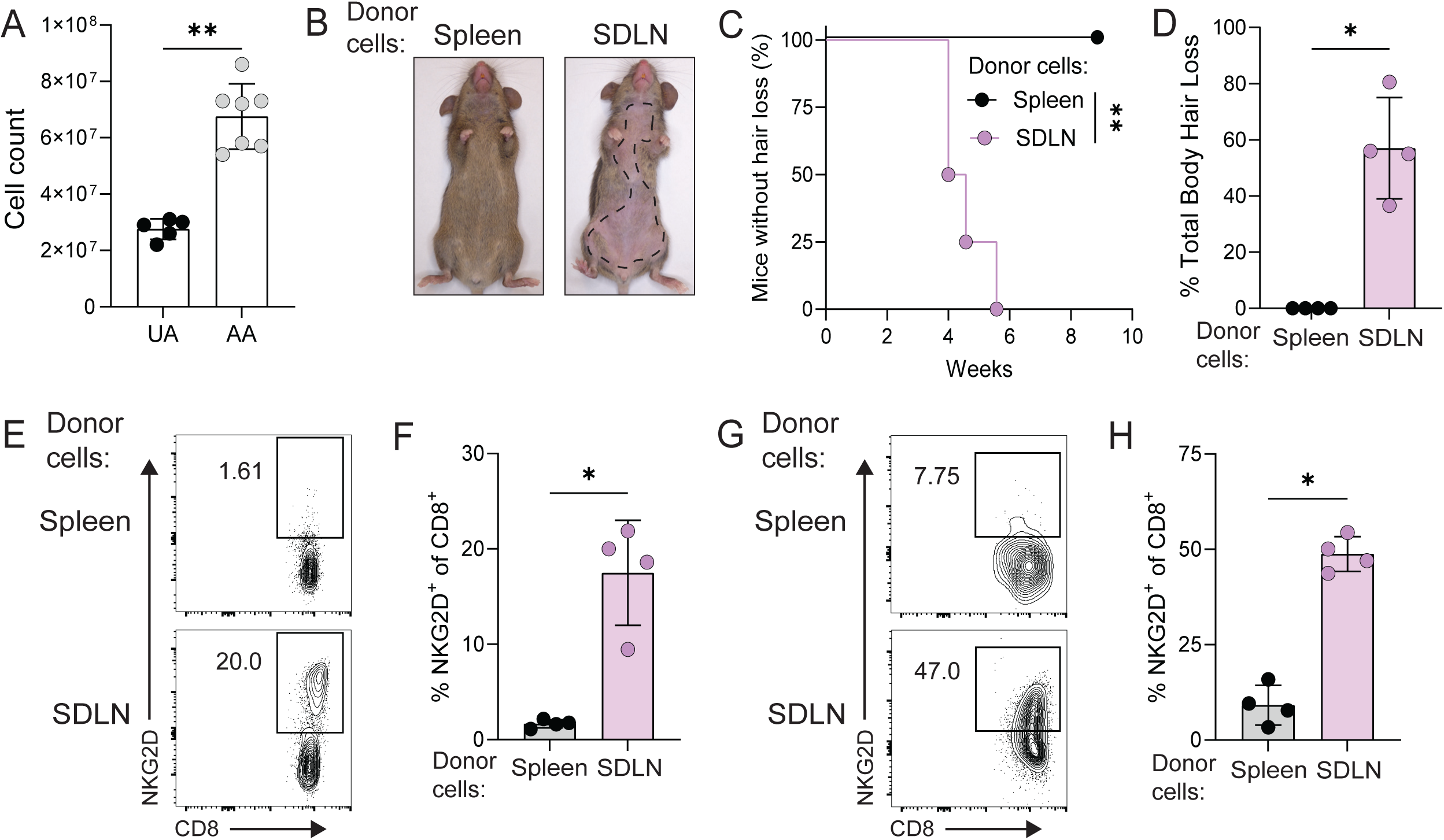
Induction of AA is restricted to the CD4 T cells in the SLDNs. **(A)** Total cell counts from spleens of UA and AA mice (mean ± SD). **p=0.0025, Mann-Whitney test. CD4 T cells were isolated from the spleens and SDLN of the same AA donor mice and transferred into recipient mice (n=4 both groups). **(B)** Representative images of ventral surface of recipient mice. **(C)** Kaplan-Meier disease curve for mice. **p=0.0062, Log-rank test. **(D)** Percent total body hair loss of mice following disease induction (mean ± SD). *p=0.0286, Mann-Whitney test. **(E)** Representative flow cytometric plots and **(F)** frequency of NKG2D^+^CD8^+^ T cells in SDLNs of recipient mice (mean ± SD). *p=0.0286, Mann-Whitney test. **(G)** Representative flow cytometric plots and **(H)** frequency of NKG2D^+^CD8^+^ T cells in skin of recipient mice (mean ± SD). *p=0.0286, Mann-Whitney test. Mice were observed two times weekly for hair loss. Data representative of two independent experiments.

Our group and others have demonstrated that SDLN-derived T cells transfer disease into recipient mice. However, it is not clear if circulating T cells are capable of transferring disease. CD4 T cells from the SDLNs and spleens were sorted, expanded *in vitro*, and transferred into recipient mice. We found that SDLN-derived CD4 T cells readily transferred disease, whereas splenic-derived CD4 T cells from the same hosts did not have the same capacity to transfer disease (Figure 2 B-D). Mice that developed AA after SDLN-derived CD4 T cell transfer exhibited the development of NKG2D^+^CD8^+^ T cells in SDLNs (Figure 2 E & F) and skin (Figure 2 G & H). These data demonstrate that pathogenic CD4 T cells capable of inducing AA are highly enriched in the regional lymph nodes

### CD4 T cells in draining lymph nodes of AA mice exhibit an enriched Th1 effector-phenotype when compared to UA mice

To gain a better understanding of the molecular mechanisms that contribute to the pathogenicity of CD4 T cells in the SDLNs, CD4^+^ T cells were isolated from the SDLNs of UA and AA mice, and RNA sequencing was performed. When evaluated through principal component analysis (PCA), we observed a distinction between the two groups of mice (Figure 3A). We identified 124 genes that were significantly up-regulated genes in the AA CD4 T cells compared to UA CD4 T cells (Figure 3B). Furthermore, we assessed our data for differences in gene expression across multiple immune-related pathways including effector molecules, transcription factors, cytokine signaling, and chemokine signaling. In CD4 T cells from AA mice, we observed an increase in expression of several genes related to activation including *Cd44, Pdcd1, Icos, and Ctla4* (Figure 3C). Interestingly, we also found several genes with increased expression relating to Th1 differentiation and function including *Stat4*, *Tbx21, Stat1, Ifng, Il18r1, Il12rb1,* and *Cxcr3* (Figure 3C). We next used the decoupleR package to infer transcription factor activity. We compared the inferred activity between UA and AA CD4 T cells (Figure 3D) and found an enriched activity in transcription factors relating to pro-inflammatory signaling including NFKB1 and RELA, as well as Th1 differentiation including TBX21 (T-bet) and SP1, a positive regulator of T-bet activity (Yu et al., 2007). Based on these findings, we next performed a gene set enrichment analysis (GSEA) of the genes with differential expression and observed a significant enrichment (enrichment score 1.81 and adjusted p (padj) value <0.001) of genes related to IFN-γ production (Figure 3E), the defining characteristic of Th1 cells. Together, these data provide evidence for an enhanced Th1 response in the CD4 T cells of AA mice.

**Figure 3.**
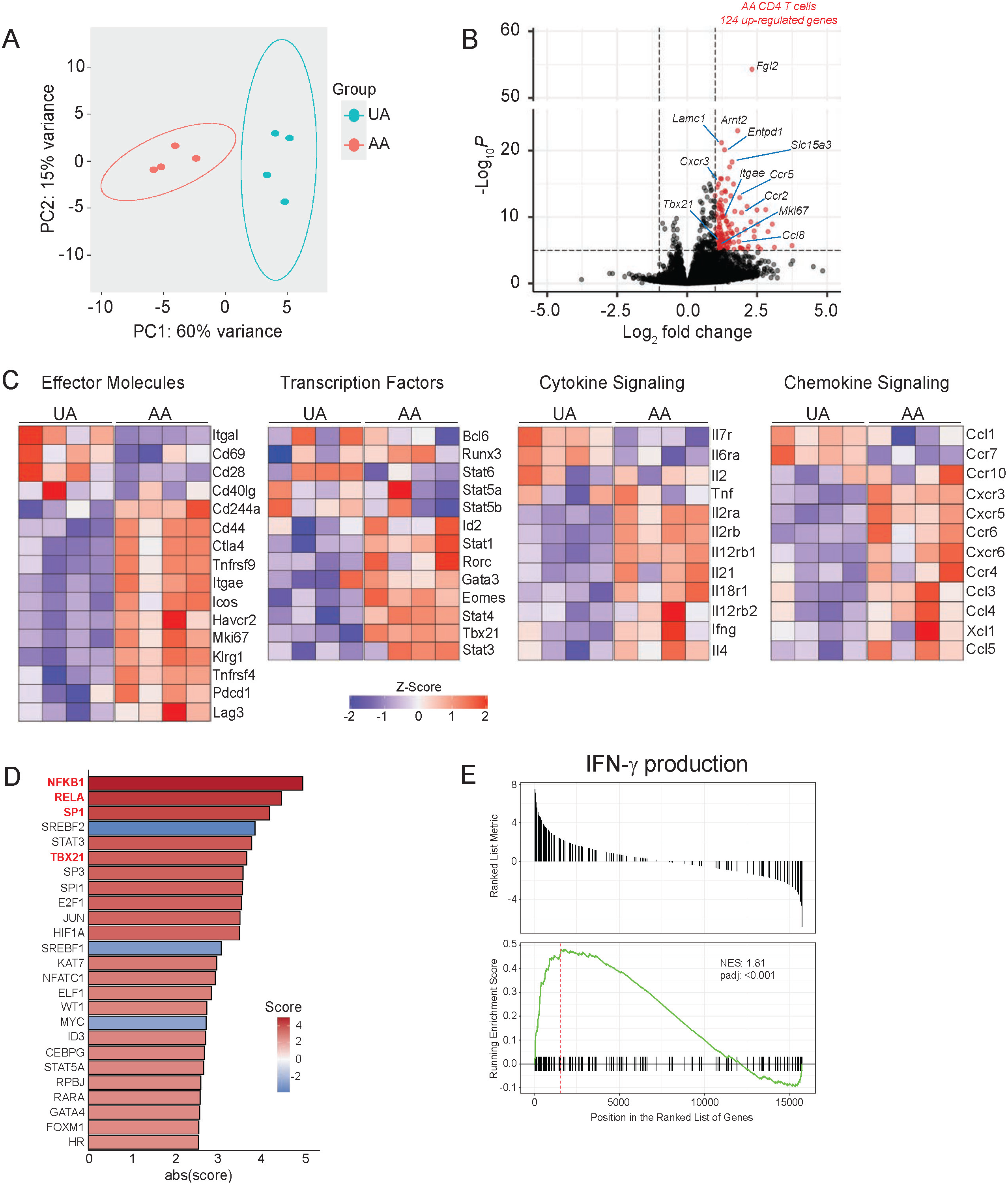
Enriched Th1 effector gene signature in CD4 T cells from AA SDLNs. Bulk CD4 T cells were FACS sorted (Live, TCRβ^+^, CD4^+^) from the SDLNs of age matched UA (n=4) and AA (n=4) mice and processed for RNA sequencing. **(A)** PCA plot of UA and AA samples **(B)** Volcano plot highlighting 124 upregulated genes in AA CD4 T cells. Log2FC>1, -Log10P>5. **(C)** Heatmaps showing differential gene expression by z-score for immune-related pathways. **(D)** Bar chart of top 25 differentially active transcription factors ranked by score. Higher activity score in AA CD4 T cells or UA CD4 T cells are indicated in red and blue, respectively. Transcription factors related to pro-inflammatory signaling and Th1 differentiation are highlighted in red. **(E)** GSEA of IFN-γ production pathway in AA CD4 T cells versus UA CD4 T cells. NES, normalized enrichment score.

Given the enhanced Th1 signature in AA CD4 T cells, we next sought to investigate the phenotypic differences between CD4 T cells of AA and UA mice. The SDLNs of AA mice exhibited an increase in the total number of CD4 T cells compared to UA mice (Figure 4A). Within the increased CD4 T cell population, we found that there were more IFN-γ producing CD4 T cells in AA mice as compared to UA mice (Figure 4B). In contrast, we found no differences in the total number of CD4 T cells producing IL-17A (Figure 4B), a T-helper type 17 (Th17) cytokine thought to play a pivotal role in other inflammatory skin diseases, such as psoriasis (Ha et al., 2014, van der Fits et al., 2009). We used spectral flow cytometry to analyze the immunophenotype of the CD4 T cell populations and performed an unsupervised clustering approach, tSNE, to compare the conventional CD4 T cell (Tconv) compartment among AA and UA mice (Figure 4C and Supplemental Figure 4). We found a minor but distinct population of cells with a greater density among the AA population (Figure 4C and 4D). Within that population, two markers of T cell activation, CD44 and CD11a, demonstrated the highest difference in expression, indicating this population has been previously activated and has likely undergone antigen-mediated stimulation (red circles) (Figure 4D). Using standard flow cytometry gating, we confirmed an increase in the expression of CD44 and CD11a among our Tconv populations in the SDLNs (Figure 4 E & F). Furthermore, these markers define a heterogenous population with varying expression of other markers of T cell activation, including CXCR3 and CD28 (Figure 4D). To investigate whether this population may be responsible for the induction of disease, we sorted this population based on distinctive markers (CD25^-^ CD44^+^CD62L^-^) and as well as a control population (CD25^-^CD44^-^CD62L^+^) of conventional CD4 T cells from the same AA mice (Figure 4G). We found that, following transfer, the effector-phenotype CD4 T cells were able to induce disease in our recipient mice whereas the control CD4 T cells were unable (Figure 4H). We found that this effector-phenotype CD4 T cell population demonstrated greater potential to produce IFN-γ following stimulation (Figure 4I). Collectively, these data indicate that the CD4 T cells of AA mice exhibit a Th1 T-effector transcriptional and phenotypic profile, and that these activated cells have the capacity to induce AA.

**Figure 4.**
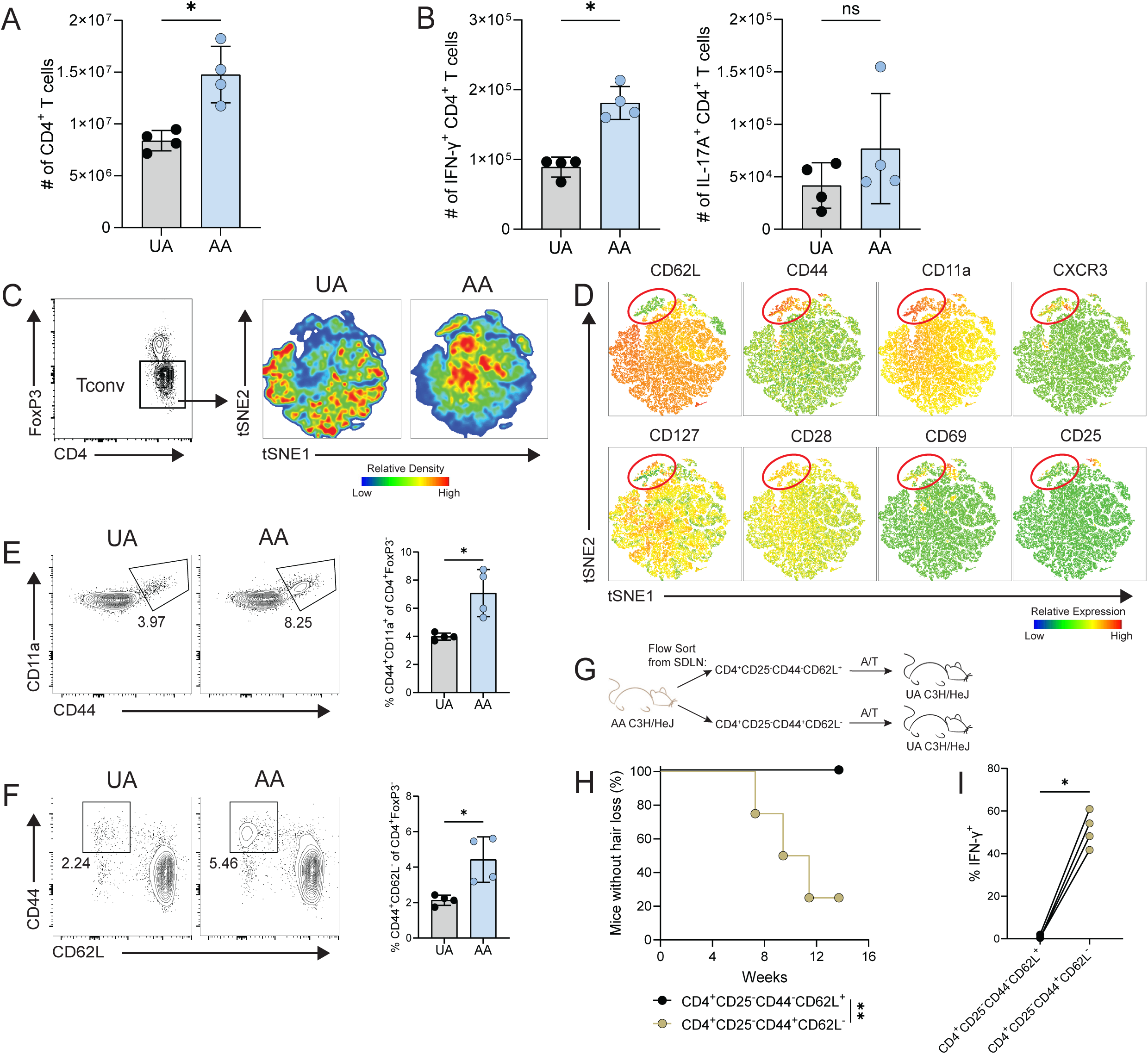
Th1 effector CD4 T cells in AA mice can induce disease. **(A & B)** SDLNs were collected from UA and AA mice and assessed for **(A)** total number of CD4 T cells and **(B)** the total number of IFN-γ^+^ and IL-17A^+^ CD4 T cells following stimulation with PMA and ionomycin (mean ± SD). *p=0.0286, Mann-Whitney test. **(C & D)** Bulk SDLN cells from the mice were collected and stained for flow cytometry. CD4^+^FoxP3^-^ conventional CD4 T cells (Tconv) were gated and down sampled to 10,000 events and concatenated. tSNE analysis was run on the concatenated sample. **(C)** tSNE analysis was separated by group (UA vs AA) showing the relative density of each group. **(D)** Heatmaps showing relative expression of different receptors within the total concatenated population. Red circle is highlighting the population of interest. **(E)** Representative flow cytometric analysis showing CD44^+^CD11a^+^ expression on Tconv CD4 T cells in the SDLNs of UA and AA mice and summary graph (mean ± SD). *p=0.0286 Mann-Whitney test. **(F)** Representative flow cytometric analysis showing CD44^+^CD62L^-^ on Tconv CD4 T cells in the SDLNs of UA and AA mice and summary graph (mean ± SD). *p=0.0286 Mann-Whitney test. **(G)** Naïve and effector Tconv CD4 T cells were FACs sorted from the SDLNs of AA mice (*naïve*: TCRβ^+^CD4^+^CD25^-^ CD44^-^CD62L^+^, *effector:* TCRβ^+^CD4^+^CD25^-^CD44^+^CD62L^-^) and underwent *in vitro* expansion for 7 days. Cells were transferred intradermally into recipient mice at approximately 1.5x10^6^ per mouse (n=8 CD4^+^CD25^-^ CD44^-^CD62L^+^, n=4 CD4^+^CD25^-^ CD44^+^CD62L^-^). **(H)** Kaplan-Meier curve for disease induction. **p=0.0040, Log-rank Test. **(I)** Frequency of IFN-γ^+^ cells were assessed prior to adoptive transfer. Each dot represents an individual experiment. *p=0.0286, Mann-Whitney test. Mice were observed two times weekly for hair loss. Data is representative of at least two independent experiments.

### CD4 T cell mediated induction of AA requires endogenous CD8 T cells

CD8 T cells have been shown to play an important role in the development of AA (Xing et al., 2014, Lee et al., 2023, Seok et al., 2023), and our data indicate that NKG2D^+^ CD8 T cell effectors emerge in mice with CD4 T cell-initiated disease. We sought to determine if CD4 T cells were required for CD8 T cell-mediated AA induction and, conversely, if CD8 T cells were required for CD4 T cell-mediated AA. To address this, we magnetically enriched CD8 T cells from AA SDLNs prior to their transfer into recipient mice. One group of recipient mice were depleted of CD4 T cells by antibody administration and compared to recipient mice that were given an isotype control antibody (Figure 5A). We found that mice treated with a CD4-targeting depleting antibody went on to develop AA similarly to those that were injected with an isotype control antibody (Figure 5 B & C). We next magnetically sorted CD4 T cells from AA SDLNs and transferred this population into recipient mice to induce AA. The endogenous CD8 T cells were depleted from one group of mice using an anti-CD8β antibody while the other group received an isotype control antibody (Figure 5D). We found that mice lacking an endogenous CD8 T cell population did not develop disease (Figure 5 E & F). These results indicate that CD4 T cells require the presence of CD8 T cells to induce the development of AA. Therefore, in CD4 T cell-mediated AA, the actions of CD4 T cells likely work upstream of, or are dependent on, the pathogenic actions of CD8 T cells.

**Figure 5.**
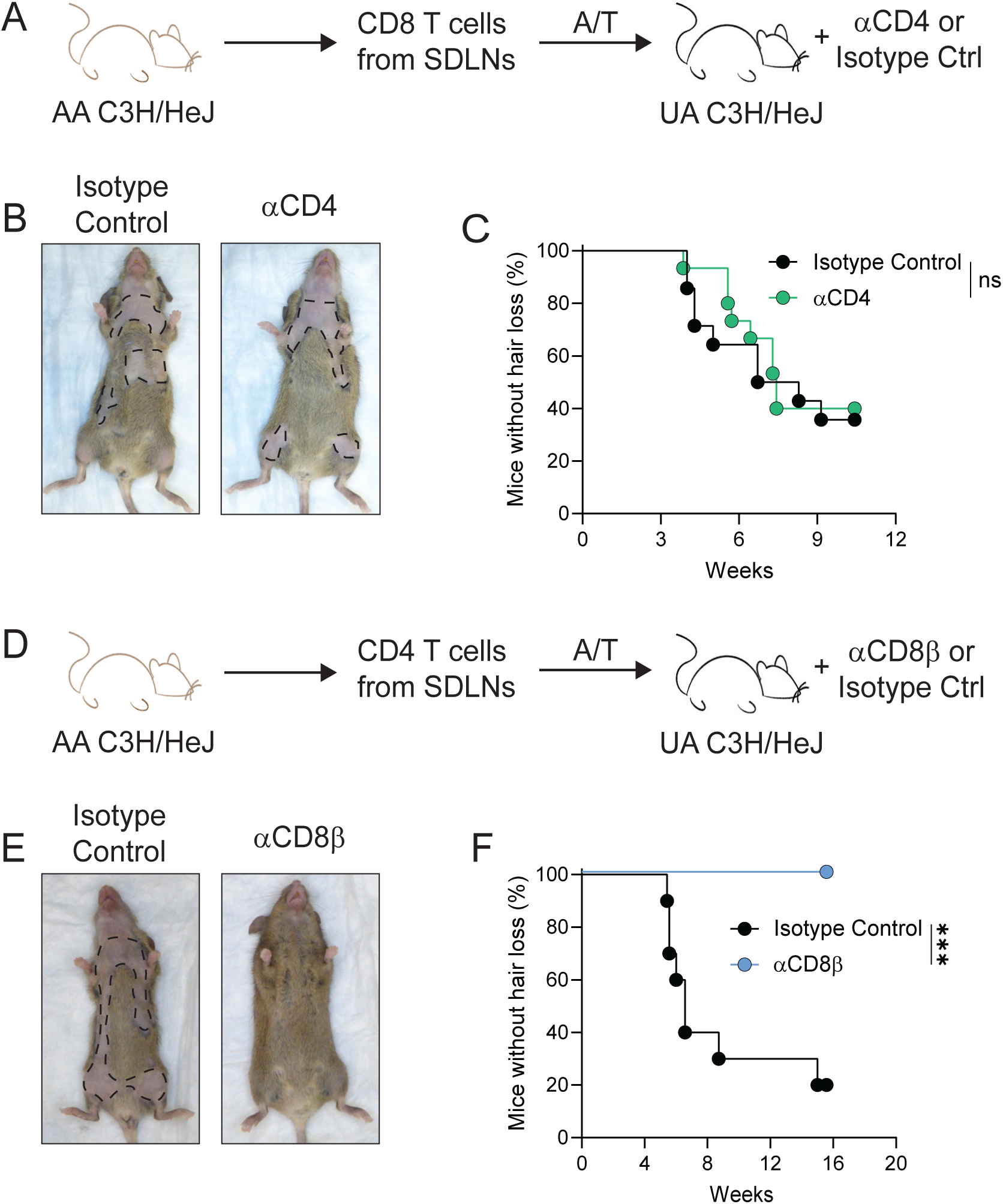
Endogenous CD8 T cells are required for CD4 T cell mediated disease induction. **(A to C)** *In vitro* activated CD8 T cells isolated from the SDLNs of AA mice were intradermally injected into recipient mice receiving 400μg intraperitoneal injections of isotype control or αCD4 antibodies (n=14 isotype control, n=15 αCD4). Mice were injected twice weekly for the duration of the experiment. **(A)** Schematic showing experimental design. **(B)** Representative images of ventral surface of mice in each group. **(C)** Kaplan-Meier disease curve for development of hair loss. ns=not significant, Log-rank test. **(D to F)** *In vitro* activated CD4 T cells isolated from the SDLNs of AA mice were intradermally injected into recipient mice receiving 400μg intraperitoneal injections of isotype control or αCD8β antibodies (n=10 each group). Mice were injected twice weekly for the duration of the experiment. **(D)** Schematic showing experimental design. **(E)** Representative images of ventral surface of mice in each group. **(F)** Kaplan-Meier disease curve showing development of hair loss. ***p=0.0003, Log-rank test. Data representative of two to three combined experiments.

### Host IFN-γ signaling is required for the induction of AA by CD4 T cells

IFN-γ is known to be important in the development of AA both by promoting MHC class I expression in the follicular epithelium and by inducing the production of chemokines that are important in the migration of T cells (Xing et al., 2014, Dai et al., 2016, Ito et al., 2004). We sought to determine whether endogenous response to IFN-γ is required for CD4 T cell-mediated AA induction. Expanded CD4 T cells from WT AA mice were transferred into either WT or IFN-γ receptor knockout (IFN-γR^-/-^) host mice. We found that IFN-γR^-/-^ mice were fully prevented from developing disease (Figure 6 A & B). We did not observe infiltration of CD8 T cells or upregulation of MHC class I throughout the hair follicles in IFN-γR^-/-^ mice (Figure 6C). Additionally, we found NKG2D-expressing CD8 T cells in the SDLNs (Figure 6 D & E) and the skin (Figure 6 F & G) of WT mice that went on to develop AA, which were largely absent in IFN-γR^-/-^ mice. Together, these data demonstrate that host IFN-γ signaling is important for the collapse of immune privilege, the development of NKG2D^+^ CD8 T cells, and the development of disease in CD4 T cell-mediated AA induction.

**Figure 6.**
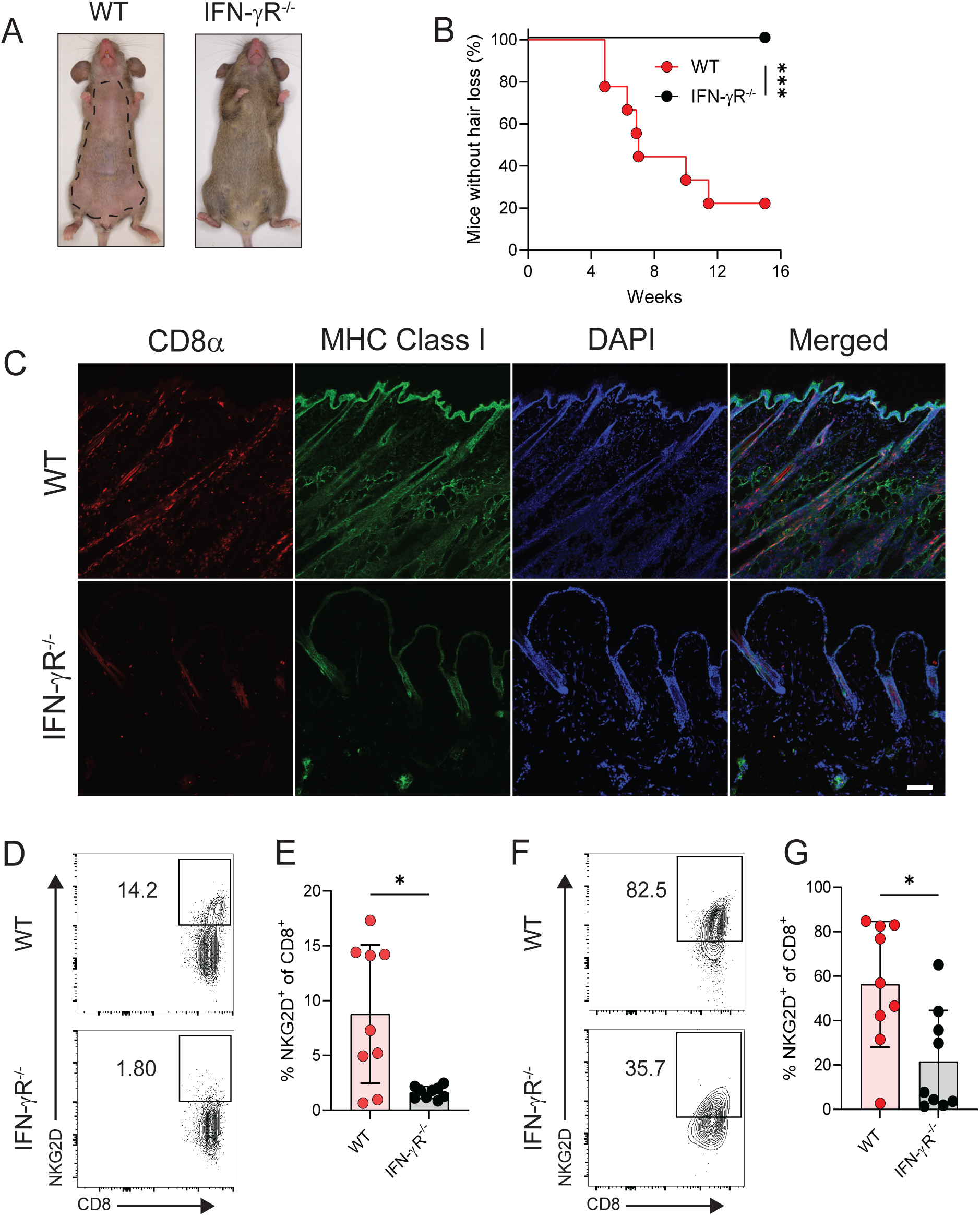
Host IFN-γ signaling is required for CD4 T cell induction of AA. CD4 T cells were isolated from the SDLNs of WT AA donor mice and underwent *in vitro* expansion. Activated CD4 T cells were intradermally injected into wild-type (WT) or IFN-γ receptor knockout (IFN-γR^-/-^) mice (n=9 both groups). **(A)** Representative ventral images of mice. **(B)** Kaplan-Meier disease curve. ***p=0.0007, Log-rank test. **(C)** Representative immunofluorescent images showing skin sections from WT and IFN-γR^-/-^ mice, stained with anti-CD8α and anti-MHC class I antibodies. Scale bar: 100μm **(D)** Representative flow cytometric analysis and **(E)** bar graph of NKG2D^+^ CD8 T cells in the SDLNs (mean ± SD). *p=0.04, Mann-Whitney test. **(F)** Representative flow cytometric analysis and **(G)** bar graph of NKG2D^+^ CD8 T cells in the skin (mean ± SD). *p=0.0188, Mann-Whitney test. Mice were observed two times weekly for hair loss. Data representative of two combined experiments.

### Effector CD4 T cells share similar transcriptional characteristics in murine and human AA

Given that the CD4 T cells of AA mice exhibit an effector Th1 phenotype, we sought to confirm this finding in human patients. We assembled and interrogated scRNA-seq data, performed an unsupervised reclustering of the T and NK cell cluster, and identified four cell populations (CD8 effector T cells, CD4 effector T cells, CD4 regulatory T cells (Treg), and NK/γδ T cells) (Figure 7A, Supplemental Figure 5) (Borcherding et al., 2020, Lee et al., 2023, Ober-Reynolds et al., 2023). We observed that all four of these populations showed an increase in the absolute cell number in AA when compared to the healthy control samples. Focusing on the CD4 effector T cell cluster, we performed differential gene expression (DEG) analysis and found an upregulation of genes associated with immune cell regulation, antigen presentation, and inflammation (Figure 7B). Interestingly, *IFNG* was found to be among the top differentially expressed genes in AA CD4 effector T cells (Figure 7B). Gene set scoring using the UCell package and two separately-generated Th1 related datasets (GSE22886 and Yasumizu et al. 2024) demonstrated an increased Th1 score among the CD4 effector T cells compared to the control samples, suggesting that AA CD4 T cells exhibit a greater Th1 signature (Figure 7C) (Yasumizu et al., 2024). Additionally, we performed a third gene set scoring analysis using a smaller, more specific list of genes that are often associated with Th1 differentiation and function. We observed a similar increase in Th1 gene scoring among the CD4 effector T cells in AA samples compared to control samples (Figure 7D). Collectively, these data support that CD4 effector cells share similar Th1 features among human samples and our mouse model.

**Figure 7.**
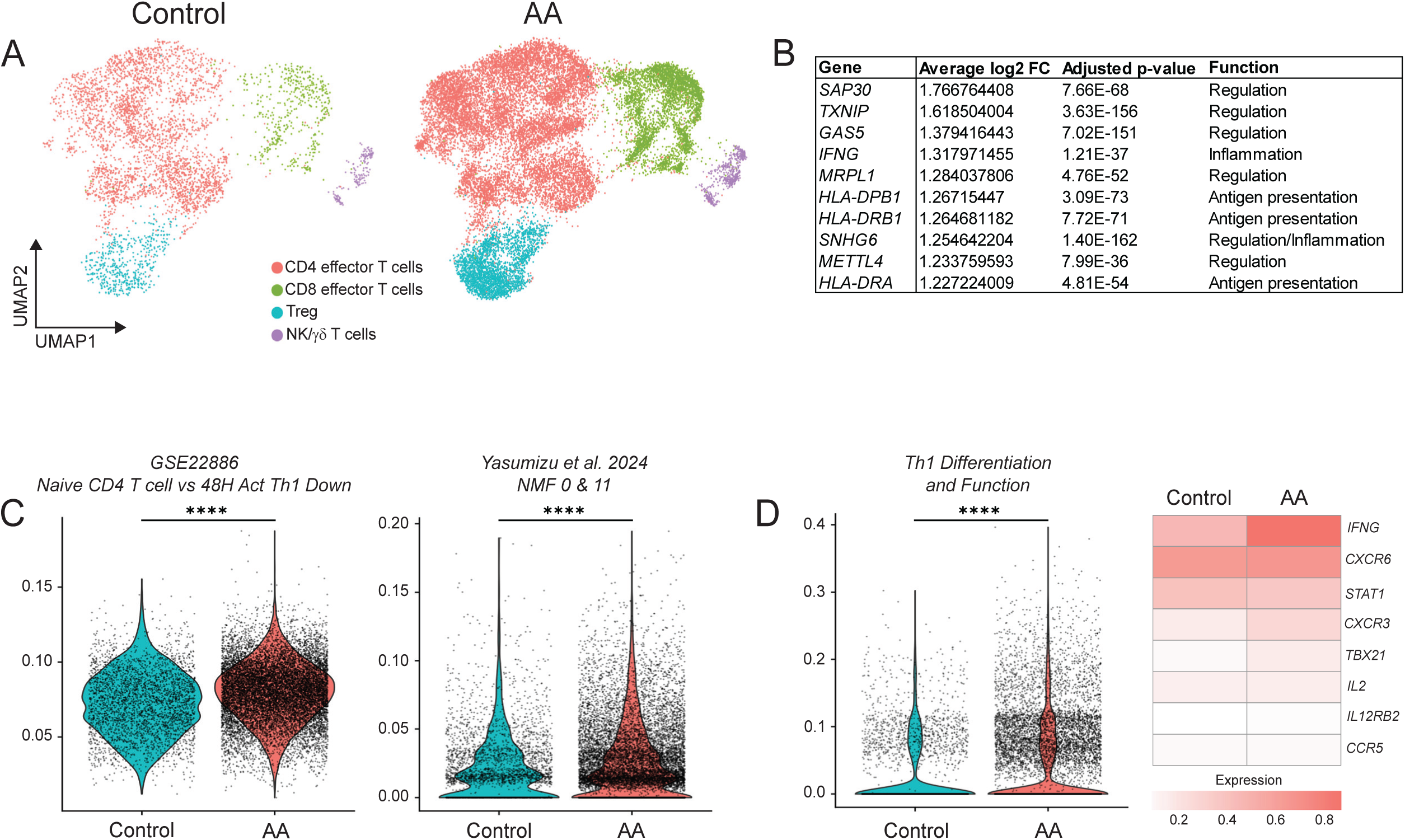
Human AA CD4 T cells express a Th1 signature. **(A)** Reclustered UMAP of the CD4 effector T cells, CD8 effector T cells, CD4 regulatory T cells (Treg) and NK/γδ T cells populations from the data in Figure 1E. **(B)** Top 10 genes up-regulated in the AA-associated CD4 effector T cells as compared to control-associated cells. Table shows upregulation of genes associated with immune cell regulation, antigen presentation, and inflammation. **(C)** Violin plots of Th1 gene set scores as calculated by UCell using MsigDB GSE22886 and dataset from Yasumizu et al. 2024. ****p<0.0001, student’s T-test. **(D)** Violin plot of Th1 gene set scores using a defined set of genes. Heatmap shows mean normalized expression for genes included in gene set. ****p<0.0001, student’s T-test.

### CD4 T cell derived IFN-γ is responsible for CD4 T cell-mediated AA development

To specifically investigate the role of IFN-γ produced from CD4 T cells in AA development, we generated CD4CreERT2-*Ifng*^flox/flox^ C3H/HeJ mice (IFN-γ^ΔCD4^), whereby IFN-γ can be efficiently deleted via a tamoxifen-inducible system (Figure 8A) (Drewry et al., 2023). AA donor mice were generated through CD8 T cell transfers to ensure that the CD4 T cell population that we would later sort from the recipient mice were made up of the genotype of the recipient. IFN-γ^ΔCD4^ and IFN-γ^WT^ (*Ifng*^flox/flox^ C3H/HeJ mice that did not harbor the *Cre* transgene) mice that developed AA were treated with tamoxifen, and loss of IFN-γ production was confirmed for CD4 T cells from IFN-γ^ΔCD4^ mice despite maintained IFN-γ production by CD8 T cells (Figure 8B). Next, CD4 T cells were magnetically sorted from the SDLNs of IFN-γ^ΔCD4^ and IFN-γ^WT^ mice with AA and expanded *in vitro*. We confirmed that IFN-γ^ΔCD4^ T cells were largely devoid of IFN-γ production (Figure 8C). To determine the importance of CD4 T cell-derived IFN-γ in AA development, we transferred IFN-γ^WT^ and IFN-γ^ΔCD4^ T cells into separate groups of recipient mice. We observed that IFN-γ^ΔCD4^ T cells were unable to induce disease in recipient mice, whereas IFN-γ^WT^ CD4 T cell recipients developed AA (Figure 8 D & E). Additionally, mice that received IFN-γ^WT^ CD4 T cells went on to develop NKG2D^+^CD8^+^ T cells in the SDLN (Figure 8F), and this population was absent in IFN-γ^ΔCD4^ CD4 T cell recipients. Together, these data demonstrate that IFN-γ produced by CD4 T cells is critical for CD4 T cell-mediated disease induction.

**Figure 8.**
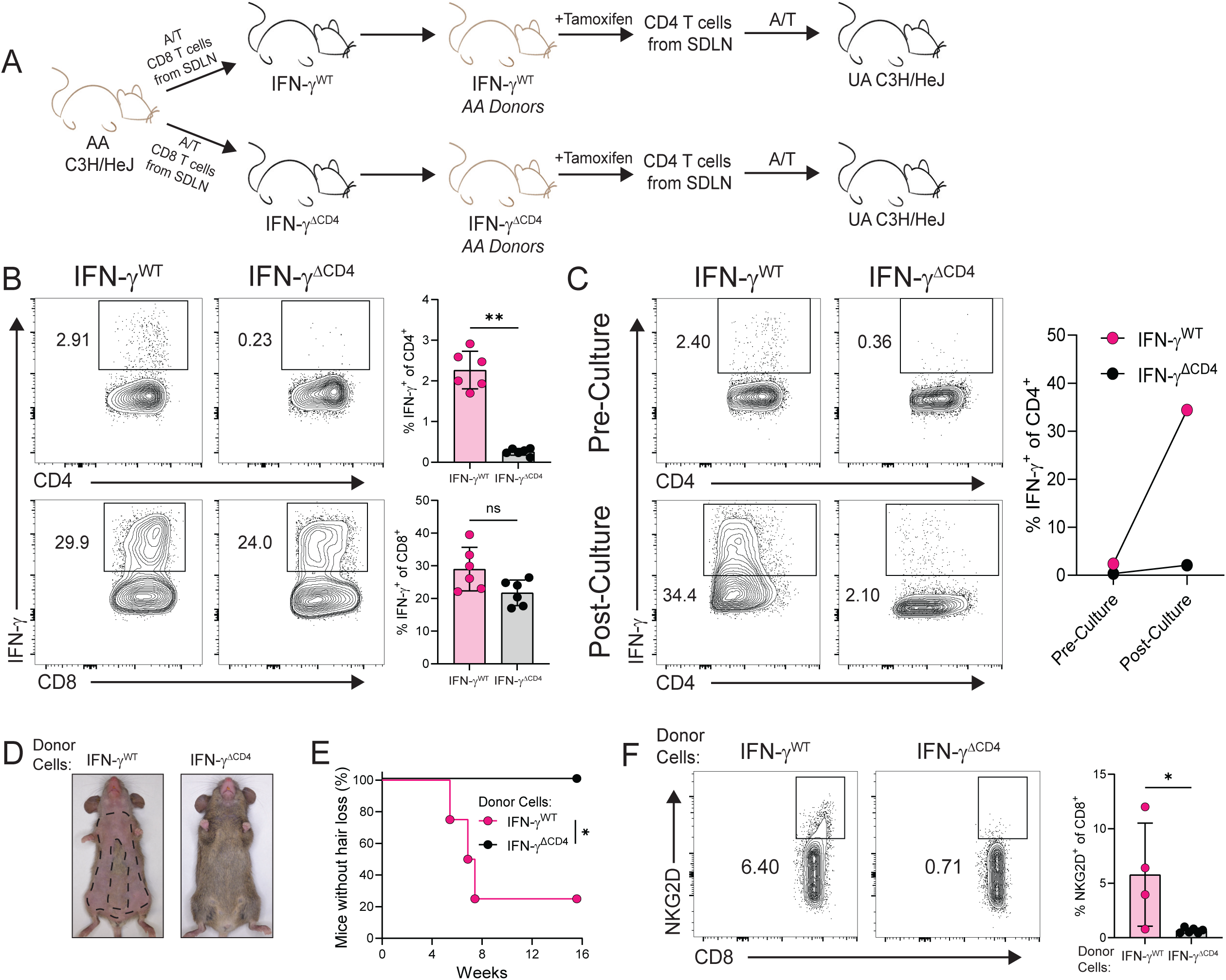
CD4 T cell derived IFN-γ is critical for the development of AA. **(A)** Experimental schematic. AA donor mice (IFN-γ^WT^ and IFN-γ^ΔCD4^) were treated with tamoxifen as indicated in the Methods. Isolated CD4 T cells from the SDLNs were pooled and activated *in vitro* and transferred into recipient mice (n=4 IFN-γ^WT^ and n= 6 IFN-γ^ΔCD4^). **(B)** Peripheral blood leukocytes were collected from mice and stimulated for 4-hours with PMA and ionomycin to assess for IFN-γ production (mean ± SD). **p=0.0022, Mann-Whitney test. **(C)** IFN-γ production from pooled SDLN CD4 T cells was assessed pre- and post-*in vitro* culture. **(D)** Representative images of ventral surface of mice in each group. **(E)** Kaplan-Meier disease curve. *p=0.0124, Log-Ranked test. **(F)** Representative flow cytometric plots and summary graphs of NKG2D^+^ CD8 T cells in the SDLNs of mice (mean ± SD). *p=0.0381, Mann-Whitney test. Data representative of two independent experiments.

## Discussion

AA is characterized as a T cell-mediated autoimmune disease of the hair follicle. Analysis of tissue samples from AA patients have demonstrated an increased presence of both CD8 and CD4 T cells around the hair follicles (Ito et al., 2008, Ito et al., 2013). Much of the previous work has primarily focused on CD8 T cells and defined them as a pivotal effector cell responsible for hair loss (Xing et al., 2014, Lee et al., 2023, Seok et al., 2023). However, AA lesions show that CD4 T cells numerically outnumber CD8 T cells (Ito et al., 2008, Ito et al., 2013, Todes-Taylor et al., 1984), yet no role for CD4 T cells have been clearly defined. Here we leveraged multiple techniques including scRNA-seq, flow cytometry, and immunohistochemistry to analyze tissue samples from AA patients. We found that T cells infiltrated the hair follicle in AA samples and made up of proportionally higher amounts of CD4 T cells compared to CD8 T cells. Further, we demonstrated that CD4 T cells from the SDLNs of AA mice are capable of inducing disease when adoptively transferred into recipient mice. We found that these cells express a Th1 effector profile with increased expression of the effector molecules CD44 and CD11a, and this ability to induce disease was dependent on the production of IFN-γ and IFN-γ signaling within the host. Together these data support a previously underappreciated role and capacity for disease induction for Th1 CD4 T cells in the development of AA.

Previous studies have shown that SDLN-derived lymphocytes from AA mice could transfer disease (McElwee et al., 2005, Wang et al., 2015). Here, we similarly show that SDLN lymphocytes from AA mice transferred disease, in contrast to SDLN lymphocytes from UA mice. Our finding that both CD4 and CD8 T cells may be involved in this model led us to further investigate the ability of CD8 and CD4 T cells to induce disease. Our data showed CD8 T cells from the SDLNs of AA mice could induce widespread disease into recipient mice, in contrast to UA CD8 T cells, a similar result to what has been shown by other groups (Xing et al., 2014, Seok et al., 2023, Dai et al., 2021). Furthermore, we demonstrated that CD4 T cells from AA mice had a robust ability to transfer disease. CD4 T cell recipient mice exhibited the emergence of NKG2D^+^CD8^+^ T cells in the SDLNs, a population of cells that is associated with disease pathogenesis (Xing et al., 2014, Petukhova et al., 2010, Seok et al., 2023). These data support that CD4 T cells have the capacity to initiate and contribute to the development of disease.

It was unclear, however, whether CD4 T cells capable of inducing AA were found in relative abundance restricted to the SDLNs or may be found systemically. Splenomegaly is a feature that has been shown in AA mice (Xing et al., 2014). We found the increased number of cells in the spleens of AA mice is made up of an increase in multiple different immune cell populations, including αβ T cells. Interestingly, within the T cell compartment, only CD4 T cells appeared to be expanded, leading us to question whether these CD4 T cells could also transfer disease. In other mouse models of peripheral autoimmunity, splenic derived T cells have been used to induce disease in the target organs, including colitis, Type 1 diabetes, and Sjögren’s disease (Ciecko et al., 2019, Ostanin et al., 2009). In our model, splenic derived CD4 T cells were not able to transfer disease similarly to SDLN-derived CD4 T cells, demonstrating that the splenic CD4 T cell pool does not harbor the same pathogenic CD4 T cell populations that are found in the lymph nodes.

Many studies have contributed to our understanding of the transcriptional and phenotypic profiles of CD8 T cells in AA, especially with regard to NKG2D-expressing CD8 T cells, which are thought to be primary effector cells in AA (Xing et al., 2014, Borcherding et al., 2020, Hashimoto et al., 2021, Lee et al., 2023, Seok et al., 2023). Fewer studies, though, have explored these profiles in the CD4 T cell population, leaving a gap in our understanding of the role of CD4 T cells in AA (Borcherding et al., 2020, Hashimoto et al., 2021, Lee et al., 2023). Using RNA-seq and spectral flow cytometry, we found that CD4 T cells in the SDLNs of AA mice exhibited a Th1 effector profile distinct from CD4 T cells from UA mice. We utilized tSNE analysis of our spectral flow cytometric data as an unbiased approach to identify markers expressed by the Tconv CD4 T cells in AA. We found a population of cells that were more represented in the AA samples, marked by the expression of CD11a and CD44, indicative of a previously activated, effector-like phenotype. We found that effector CD4 T cells (CD25^-^ CD44^+^CD62L^-^) were potent producers of IFN-γ and transferred disease.

A variety of disease models indicate that CD4 T cell helper functions are important for a robust and memory-generating CD8 T cell response. In a model using glia-tropic mouse hepatitis virus, depletion of CD4 T cells early in the immune response compromised the CD8 T cell function and decreased viral clearance (Phares et al., 2012). Additionally, other models of bacterial or viral infections have demonstrated that during CD8 T cell priming, the absence of CD4 T cell help results in an inadequate expansion of the effector CD8 T cells either during the primary or subsequent challenges (Sun and Bevan, 2003, Shedlock and Shen, 2003, Janssen et al., 2003, Novy et al., 2007, Kumamoto et al., 2011). These data support that CD4 T cell help is important in the early activation stages of a CD8 T cell response, which could be in part by licensing of dendritic cells through the CD40-CD40L pathway (Schoenberger et al., 1998, Ballesteros-Tato et al., 2013, Bennett et al., 1998, Ferris et al., 2020). Previously, using scRNA-seq, our group found that antigen presenting cells in AA skin showed an increase gene signature for CD40 signaling, indicating one potential mechanism by how CD4 T cells could be contributing to AA (Borcherding et al., 2020). Together these data led us to question how CD4 and CD8 T cells cooperate in AA and what role CD4 T cells may have in helping in the induction of NKG2D expressing CD8 T cells. We found CD8 T cells can transfer disease whether or not CD4 T cells are present. However, CD4 T cells lacked the ability to transfer disease if CD8 T cells were not present. This suggests that, in AA development, CD4 T cells may have a role in the priming of CD8 T cell responses that are needed in order for disease to emerge. A prior study showed that in patients in an acute phase of disease, reported as less than six months duration and between 1 to 20 lesions, there were more CD4 T cells present in the skin compared to CD8 T cells, providing another example of how CD4 T cells may be more important during early stages of disease (Ito et al., 2013). A more recent study showed that in the skin graft model of AA, if CD4 T cells are depleted, there was a significant delay in AA induction (Lee et al., 2023, McElwee et al., 1998). Furthermore, prior studies have indicated that depletion of CD8 T cells protects AA skin graft recipients from developing disease (Lee et al., 2023). Together these data invite future studies to help dissect what specific mechanisms CD4 T cells use to help promote CD8 T cells in AA, potentially including the CD40-CD40L pathway.

IFN-γ has been implicated as a driver of the collapse of IP of the hair follicle and a critical cytokine in AA pathogenesis (Ito et al., 2004, Xing et al., 2014). Furthermore, IFN-γ stimulation of the hair follicles induced the expression of the pro-inflammatory chemokines CXCL9 and CXCL10, which have been shown to be important in the migration of T cells to the hair follicle and in AA development (Dai et al., 2016). Previous studies have specifically targeted IFN-γ to investigate the role in AA induction; IFN-γ deficient mice were protected from developing disease using the skin graft model of AA (Freyschmidt-Paul et al., 2006), and, in another study, antibody mediated neutralization of IFN-γ blocked the AA induction in AA skin graft recipient mice (Xing et al., 2014). We investigated the role of IFN-γ signaling in mice following CD4 T cell mediated induction using mice exhibiting genetic ablation of the IFN-γR. Following the transfer of wild-type CD4 T cells, IFN-γR^-/-^ mice were protected from developing AA. Our data demonstrate that responsiveness to IFN-γ is critical to disease development following CD4 T cell-mediated AA induction. Taken together, these data implicate IFN-γ as an essential cytokine in AA pathogenesis in a variety of models. It is still unclear, however, which cell populations are responding to IFN-γ. The hair follicle expresses the IFN-γ receptor and following stimulation with IFN-γ, upregulates MHC class I molecules, a hallmark of the IP collapse seen in AA (Ito et al., 2004, Ito et al., 2005, Paus and Bertolini, 2013). Furthermore, CD8 T cells express the IFN-γ receptor and signaling through this pathway is important for a variety of cellular functions. Following lymphocytic choriomeningitis viral infection, IFN-γ receptor deficient CD8 T cells showed a reduced expansion as compared to IFN-γ receptor sufficient cells (Whitmire et al., 2005). In another study of graft rejection, using intravital imaging, CD8 T cells stimulated with IFN-γ moved faster through the skin and traveled a greater distance (Bhat et al., 2017). Additionally, these cells were also more cytotoxic when challenged with keratinocytes loaded with relevant peptides. These data support that the hair follicle, CD8 T cells, or both may be responding to IFN-γ in AA to engender disease development.

Using our scRNA-seq data, we more closely investigated the CD4 effector T cell population that is present in the skin of human patients. Our data indicate CD4 effector T cells are more highly represented in AA samples when compared to healthy control samples, and these cells express a Th1 signature. We found *IFNG* as one of the top differentially expressed genes in CD4 T cells from AA lesions when compared with T cells from healthy controls. A recent study of murine skin found that the *Ifng* gene is also among the top differentially expressed genes in AA CD4 T cells (Lee et al., 2023). This data along with our mouse RNA-seq data points to IFN-γ potentially being a critical part of CD4 T cells involvement in AA. Th1 cells have often been associated with development of other autoimmune diseases and models, including Sjögren’s disease, colitis, and experimental autoimmune encephalomyelitis (Wang et al., 2024, Domingues et al., 2010, Harbour et al., 2015). In a recent study of Sjögren’s disease, deletion of IFN-γ production from CD4 T cells eliminated their ability to induce glandular disease (Wang et al., 2024). In AA, we hypothesized that CD4 T cell-derived IFN-γ may be an essential function of CD4 T cells in inducing disease. Deleting IFN-γ production from CD4 T cells eliminated their ability to transfer disease. This data is supported in part by our earlier findings (Figure 4I), indicating that effector T cells produced greater amounts of IFN-γ compared to their naïve counterparts. Together, this data supports that Th1 cells can contribute to the development of AA.

Overall, our study describes a critical role for CD4 T cells in AA. We found that CD4 T cells from AA mice have a distinct Th1 transcriptional and phenotypic profile, which highlights a specific population of cells that can induce disease. These data support further investigation into the CD4 T cells in human patients to better understand the characteristics that may help define populations important in disease pathogenesis. Additionally, our data demonstrate the potential of CD4 T cells to be important in the early stages of disease development, prompting future studies to better understand the kinetics of CD4 T cells in AA and define potential mechanisms that may be useful for therapeutic intervention.

## Materials and Methods

### Mice

All experiments used female C3H/HeJ mice purchased from The Jackson Laboratory (Stock #000659). Mice were kept in pathogen-free conditions at the University of Iowa Animal Facility. IFN-γR^-/-^ mice on the C3HeB/FeJ background were a generous gift from the Goverman lab (Department of Immunology, University of Washington) and were backcrossed for seven generations onto the C3H/HeJ background (Simmons et al., 2014). B6(129X1)-Tg(Cd4-cre/ERT2)11Gnri/J (CD4CreERT2) mice were purchased from The Jackson Laboratory (Stock #022356) and backcrossed onto the C3H/HeJ background for at least 11 generations. *Ifng*^flox/flox^ mice on the C57BL/6 background were a generous gift from the Harty Lab (Department of Pathology, University of Iowa) and were backcrossed for 10 generations onto the C3H/HeJ background (Drewry et al., 2023). CD4CreERT2 and *Ifng*^flox/flox^ mice were bred to produce CD4CreERT2-*Ifng*^flox/flox^ mice, and Cre genotypes were confirmed by PCR. All animal procedures were approved by the University of Iowa Institutional Animal Care and Use Committee (IACUC) and the Department of Veteran’s Affairs IACUC.

### Study Participants

Healthy control patients and patients with AA were recruited from the Department of Dermatology at University of Iowa Health Care, and written informed consent was obtained in compliance with the guidelines of the institutional review board-approved protocol #201706728 and the Declaration of Helsinki.

### T cell isolation for *in vitro* culture and induction of AA

SDLNs (inguinal, brachial, axillary, and cervical) and whole spleens were isolated from UA and/or AA affected mice. Single-cell suspensions of cells were made by mechanically dissociating tissue through a 70μm cell strainer. Splenocytes were depleted of erythrocytes by RBC Lysis Buffer (Thermo Fisher Scientific, 00-4333-57).

From the single cell suspensions, CD4 or CD8 T cells were isolated using magnetic bead separation per manufacturer instructions (MACS, Miltenyi Biotec, CD4 (L3T4) 130-117-043, CD8α (Ly-2) 130-117-044). Enriched T cells were then placed into culture in complete RPMI 1640 media containing 10% fetal bovine serum (FBS) (Thermo Fisher Scientific), 2mM L-glutamine (Thermo Fisher Scientific), 10mM HEPES (Thermo Fisher Scientific), 1x antibiotic-antimycotic (100U/mL penicillin, 100μg/mL streptomycin, and 250ng/mL Amphotericin B) (Thermo Fisher Scientific), 1mM Na Pyruvate (Thermo Fisher Scientific), and 50μm 2-mercaptoethanol (Avantor). Culture media was supplemented with IL-2 (30U/mL), IL-7 (12.5ng/mL), IL-15 (25ng/mL) (Cytek, Biolegend, Cell Signaling, and Thermo Fisher Scientific), and mouse T-activator CD3/CD28 Dynabeads (Thermo Fisher Scientific). Cells were activated and expanded for 6-8 days at 37°C and 5% CO2. In some experiments, bulk SDLN cells were placed into culture in complete RPMI media containing 10% FBS, 2mM glutaMAX (Thermo Fisher Scientific), and 100U/mL penicillin streptomycin (Thermo Fisher Scientific). Cells were expanded for 6-8 days followed by magnetic enrichment of CD4 T cells by MACs separation. Purity of T cells was assessed post culture and cultures with greater than 98% purity were used.

To induce AA in C3H/HeJ mice, cells were collected and CD3/CD28 Dynabeads were magnetically separated. The cells were then washed in sterile PBS and resuspended to the desired concentration in sterile saline or PBS. Cells were transferred intradermally into 10–14-week-old normal hairy C3H/HeJ mice (Wang et al., 2015). For experiments using CD8 T cells, cells were injected between 5-10x10^6^ per mouse and for experiments using bulk CD4 T cells, cells were injected between 3.33-30x10^6^ per mouse. For experiments using naïve and effector CD4 T cells, cells were injected at 1.5x10^6^ per mouse. Mice were checked twice weekly for the development of AA on the ventral surface. Hair loss was quantified using Fiji software by calculating % total body hair loss and subtracting % areas of partial or total body hair loss.

### *In vivo* depleting antibodies

For *in vivo* depletion studies, antibody treatments were started one day prior to the transfer of lymphocytes. Anti-CD4 (BioXCell, BE0003), anti-CD8β (Lyt 3.2, BioXCell, BE0223), or isotype control IgG (BioXCell) were administered by intraperitoneal injection (400μg) two times weekly for the duration of the experiments.

### Preparation of tissue cell suspensions

SDLNs or whole spleens were mechanically dissociated and filtered through a 70μm cell strainer. Splenocytes were depleted of erythrocytes by RBC Lysis Buffer (Thermo Fisher Scientific) and washed in PBS. To prepare skin single cell suspension, hair was removed from the mice using hair clippers. Skin was collected, defatted, and finely minced using scissors and digested for 60 minutes at 37°C with collagenase IV (1.5mg/mL, Worthington) and deoxyribonuclease I (DNase I) (0.025mg/mL, Roche) in RPMI 1640 with 3% FBS in a shaker. Digested skin was filtered through a 70μm cell strainer and washed twice with cold RPMI 1640 with 10% FBS before staining.

Human skin samples were collected, defatted, and finely minced using scissors and digested for 60 minutes at 37°C with Liberase TL (350μg/mL, Sigma-Aldrich) and DNase I (75μg/mL, Worthington). Digested samples were then mechanically dissociated and filtered through a 70μm cell strainer and washed twice with PBS containing 2% FBS before staining.

### Flow cytometry and fluorescence activated cell sorting

Single cell suspensions were stained with Live Dead Blue or Zombie UV (Thermo Fisher Scientific and Biolegend) for 30 minutes at 4°C prior to surface markers. Surface proteins were then labelled with fluorophore-conjugated antibodies (Tables 1 & 2) for 20-30 minutes at 4°C in PBS with 2% FBS. In some experiments, Ghost Dye antibodies (Cytek Biosciences) were included into the surface marker cocktail to detect viable cells. For intracellular staining of transcription factors and/or cytokines, cells were fixed with FoxP3/Transcription Factor Staining Buffer Kit (Cytek Biosciences, TNB-0607) according to manufacturer’s instructions and then incubated with fluorophore-conjugated antibodies. For cytokine staining, cells were stimulated for 4-5 hours with phorbol 12-myristate 13-acetate (PMA) and ionomycin (Biolegend, 423302) with Brefeldin A (Biolegend, 420601) and washed two times in PBS with 2% FBS before staining. Flow cytometry data were acquired using a BD LSRII (BD Biosciences) or Cytek Aurora (Cytek Biosciences) and analyzed with FlowJo V10 software (Treestar).

For cell sorting, cells were incubated with fluorophore-conjugated antibodies in PBS with 2% FBS for 30 minutes at 4°C. The samples were then washed and resuspended in PBS supplemented with 2% FBS and 10mM HEPES (Thermo Fisher Scientific). Cells were sorted on either the BD Aria Fusions sorter (Becton Dickinson) or the Cytek Aurora CS (Cytek Biosciences). Viable cells were gated on forward and side scatter and Hoechst 33458 staining.

FlowJo software was used for tSNE analysis. For the analysis, cells were gated on forward and side scatter (FSC-A/SSC-A), single cells (FSC-H/FSC-A), live, CD3^+^, CD4^+^, and FoxP3^-^ to get conventional CD4 T cells (Tconv) (Supplemental Figure 4). Tconv cells were down sampled to 10,000 events and concatenated from all samples to perform tSNE analysis (Linderman et al., 2019, Belkina et al., 2019), and then separated by disease state. Heatmaps of individual markers were assessed on the total concatenated sample.

### Immunostaining

Mouse skin was embedded in optimal cutting temperature (OCT) compound and 30μm sections were cut. Sections were fixed in cold (-20°C) acetone for 10 minutes, washed with wash buffer (1x TBS containing Ca and 0.09% NaN_3_) for 10 minutes, and blocked with wash buffer containing 5% bovine serum albumin (RPI), and 5% goat serum (Sigma-Aldrich) for 60 minutes at 37°C. After blocking, the sections were stained with rat anti-CD8α (Clone 53-6.7, Thermo Fisher Scientific, 14-0081-85) and biotin mouse anti-H2K^k^ (Clone 36-7-5, Biolegend, 114903) overnight at 4°C at a 1:100 dilution in blocking buffer. Following the primary antibody, sections were stained with fluorescently tagged secondary antibodies, anti-rat Alexa Fluor 568 (1:600, Thermo Fisher Scientific, A-11077) and Alexa Fluor 488 anti-biotin (1:400, Jackson Immunoresearch, 200-542-211). For detection of nuclei, sections were stained using 1μg/mL DAPI (Cell Signaling, 4083) for 5 minutes at room temperature. Antifade mountant with DAPI (Thermo Fisher Scientific, P36962) was used as the mounting medium. Immunofluorescent images were captured on a Zeiss LSM 880 confocal microscope and images were compiled using Fiji.

Human scalp tissue was fixed in 10% neutral buffered formalin and embedded in paraffin. Paraffin sections were cut at 5μm and CD3 staining was performed by the University of Iowa Comparative Pathology Laboratory. Images were captured using an Olympus BX63 microscope.

### Tamoxifen Treatment

Tamoxifen (Sigma-Aldrich, T5648) was suspended in corn oil and 10% ethanol and treatments were performed by intraperitoneal injection. Three to four doses of 4mg were administered every other day into Cre positive (IFN-γ^ΔCD4^) and Cre negative (IFN-γ^WT^) mice. Excision of IFN-γ in the CD4 T cells was confirmed by intracellular cytokine staining of peripheral blood after PMA/Ionomycin stimulation prior to collection of SDLNs. Production of IFN-γ from CD8 T cells was also examined to confirm CD4 T cell specific deletion. Deletion of IFN-γ was also confirmed in SDLNs before and after the *in vitro* expansion of the CD4 T cells.

### Bulk RNA Sequencing

Mice were induced to develop AA by full thickness skin grafts (McElwee et al., 1998) and allowed to develop disease. For cell isolation, SDLNs were removed from aged-matched AA and UA mice. Single cell suspensions were made and live TCRβ^+^CD4^+^ T cells were sorted using a FACS Aria Fusion (Becton Dickinson). Cells were washed and frozen prior to RNA isolation. Libraries were generated using the Truseq Stranded mRNA kit (Illumina 20020595), following the manufacturer’s protocol with 100ng RNA input. Libraries were analyzed for quality using qubit (ThermoFisher, Q32854) and size distribution using Agilent Tapestation. Samples were pooled and sequenced on Novaseq S4 200 cycle flowcell (Illumina) according to the manufacturer’s recommendations to achieve 20 million paired reads per sample at MedGenome Inc. Reads were aligned to the reference genome using the STAR aligner (version 2.7.10a) and a genome index was generated using the GRCm39 primary assembly (FASTA) and the corresponding GENCODE vM30 annotation (GTF). Gene-level counts were quantified during alignment using theL--quantMode GeneCountsLoption in STAR. Gene counts were imported into R (version 4.4) and consolidated into a single expression matrix for all eight samples. Lowly expressed genes were filtered out by retaining genes with a total count greater than 30 across all samples and fewer than 6 zero-count samples. Transcript-to-gene mappings were performed using the GENCODE vM30 annotation file (tx2gene.gencode.vM30.csv), and gene-level counts were aggregated by summing counts across transcript isoforms. Differential expression analysis was performed using the DESeq2 package (version 1.46.0). A DESeqDataSet object was created from the filtered count matrix and sample metadata, and theLDESeqLfunction was applied to normalize counts and fit a negative binomial model. Variance-stabilizing transformation (VST) was applied to the normalized counts for visualization and downstream analyses. Principal component analysis (PCA) was performed on the VST-transformed counts using the top 500 most variable genes, and 95% confidence ellipses were added to visualize group separation using theLggplot2Lpackage. A volcano plot was generated to visualize differential gene expression (DEG) using theLEnhancedVolcanoLpackage. Genes with significant differential expression (adjusted p-value < -log_10_(5) and |log2FC| > 1) were highlighted in red. Heatmaps were generated for selected marker genes and grouped by functional categories. Expression levels for each gene were extracted from the VST-transformed matrix, centered, and scaled across all samples. Rows and columns were clustered based on the scaled expression values (z-score), and boxes were colored according to this z-score. Plots were generated using theLggplot2Lpackage. Transcription factor (TF) activity was inferred using theLdecoupleRLpackage and the CollecTRI network. Significant differentially expressed genes (adjusted p-value < 0.05) between the AA and UA were used as input, with theLstatLvalue from DESeq2 differential expression analysis serving as the weight. The top 25 differentially inferred TFs were extracted for visualization. Gene set enrichment analysis (GSEA) was performed using the clusterProfiler package on the differentially expressed genes. The Gene Ontology database GO:0032609 (IFN-γ production) was used for analysis.

### Human scRNA-seq data analysis

Previously published human scRNA-seq data were acquired from the GEO database using accessions GSE212450 (Ober-Reynolds et al., 2023), GSE145095 (Borcherding et al., 2020), and GSE233906 (Lee et al., 2023). For each dataset, the filtered barcode, feature, and matrix files were downloaded and loaded into R. scDblFinder was used to remove doublets from each sample before being merged into a single Seurat object (min.cells = 5, min.features = 500). This Seurat object was filtered to include percent.mt < 20, nCount_RNA < 60000, nCount_RNA > 800, and db.class == “singlet”. Variable Features were identified using default parameters and any mitochondrial, ribosomal, cell cycle, TCR, BCR, and other select genes that have been noted to cause issue were removed. The data was normalized by log-transformation and scaled before being integrated using PCA and the Harmony package. The harmonized data was used to generate a uniform manifold approximation map (UMAP) using the FindNeighbors, FindClusters, and RunUMAP functions. The UMAP was generated using a resolution of 0.3 and clusters were labeled based on the expression profiles and DEG results. Differential abundance testing was performed using the miloR package. Three samples were excluded from this analysis due to immune cell flow sorting or poor cell acquisition (>90% keratinocyte abundance). The remaining samples were converted into a milo object and applied to the harmonized data. A graph was built, and neighborhoods were created using k = 20, d = 50, prop = 0.1, and refined = T. Differential testing was done using disease state (AA versus control) and the results were overlayed on the existing UMAP. The logFC abundances were colored for all neighborhoods (red = AA, blue = control) below a false discovery rate (FDR) of 10%. The neighborhoods were then annotated according to the most abundant cell type in each neighborhood, and the differential abundance testing results were plotted as a beeswarm plot, grouped by the cell types identified (FDR 10%). To analyze the T cells, lymphoid cells were isolated from the dataset, processed similarly to the full dataset, and labeled according to the major subsets identified. The CD4 effector T cell population was further isolated and DEG analysis was performed on this population, comparing AA versus control cells. Genes with a pct.1 greater than 0.1 and adjusted p-value less than or equal to 0.05 were filtered out and the data was sorted by log2 fold-change (FC) and presented as the top 10 highest log2 FC. The UCell package was used to score the normalized expression of the CD4 effector T cell populations using different Th1 gene signature lists. Two of the gene signature lists were collected from (1) the GEO database using accession # GSE22886, and (2) two non-negative matrix factorization (NMF) populations expressing Th1 gene signatures, NMF 0 and 11, from a prior study (Yasumizu et al., 2024). A selective gene list was generated using genes that have been used to define Th1 cells in Figure 7D. The UCell scores for each of the three gene signature lists were used to generate violin plots. A heatmap was made of the selective gene list showing the average expression of each gene compared between AA and control CD4 effector T cells colored according to the absolute expression level.

### Statistical Analysis

For biological (non-omics) analysis, data were analyzed using GraphPad Prism (version 10) software. Two groups were compared using an unpaired nonparametric t-test (Mann-Whitney test) or a paired nonparametric t-test (Wilcoxon signed rank test). Kruskal-Wallis test with Dunn’s multiple comparisons test was used for mean differences comparison from multiple groups. Log-rank (Mantel-Cox) tests were used to analyze the hair loss curve. Data is presented as means ± SD. Statistical significance has been indicated within the figure legends with asterisks. All experiments were repeated at least two times. All bioinformatic analyses were performed using R. For the scRNA-seq data, a student’s t-test was used to compare the CD4 effector T cells in AA versus control samples in Figure 7.

## Supporting information

Table 1 - Mouse Flow Antibodies

Table 2 - Human Flow Antibodies

## Data Availability

All RNAseq data for SDLN CD4^+^ T cells have been deposited in NCBI’s Gene Expression Omnibus and are accessible through GEO accession number GSE295313.

## Acknowledgements

This work was supported by National Institutes of Health grants K08AR069111 (A.J.), R01AR077194 (A.J.), T32AI007485 (S.J.C.) and the Department of Veteran’s Affairs grant 5I01BX004907 (A.J., Biomedical Laboratory Research and Development)

The data presented herein were obtained at the Flow Cytometry Facility, which is a Carver College of Medicine / Holden Comprehensive Cancer Center core research facility at the University of Iowa. The facility is funded through user fees and the generous financial support of the Carver College of Medicine, Holden Comprehensive Cancer Center, and Iowa City Veteran’s Administration Medical Center. Research reported in this publication was supported by the National Center for Research Resources of the National Institutes of Health under Award Number 1 S10 OD034193-01. The authors would like to acknowledge use of the University of Iowa Central Microscopy Research Facility, a core resource supported by the University of Iowa Vice President for Research, and the Carver College of Medicine.

