## Supplemental Figures for "Th1 effector CD4 T cells rely on IFN-γ production to induce alopecia areata"

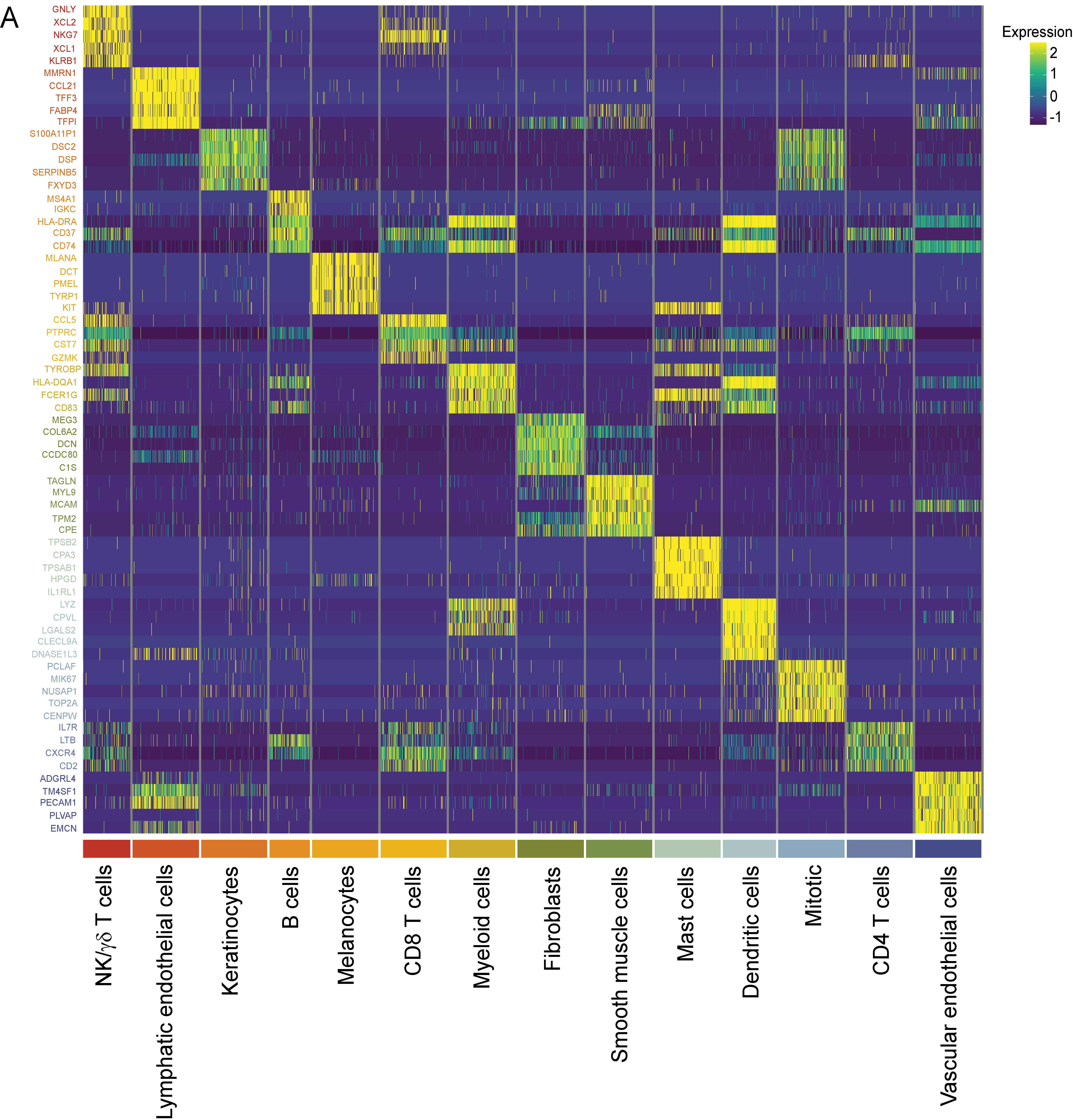

**Supplemental Figure 1. Human scRNA-seq cell type identification. (A)** Heatmap of the top differentially expressed genes, ranked by percent difference, used to identify each of the 14 cell populations.

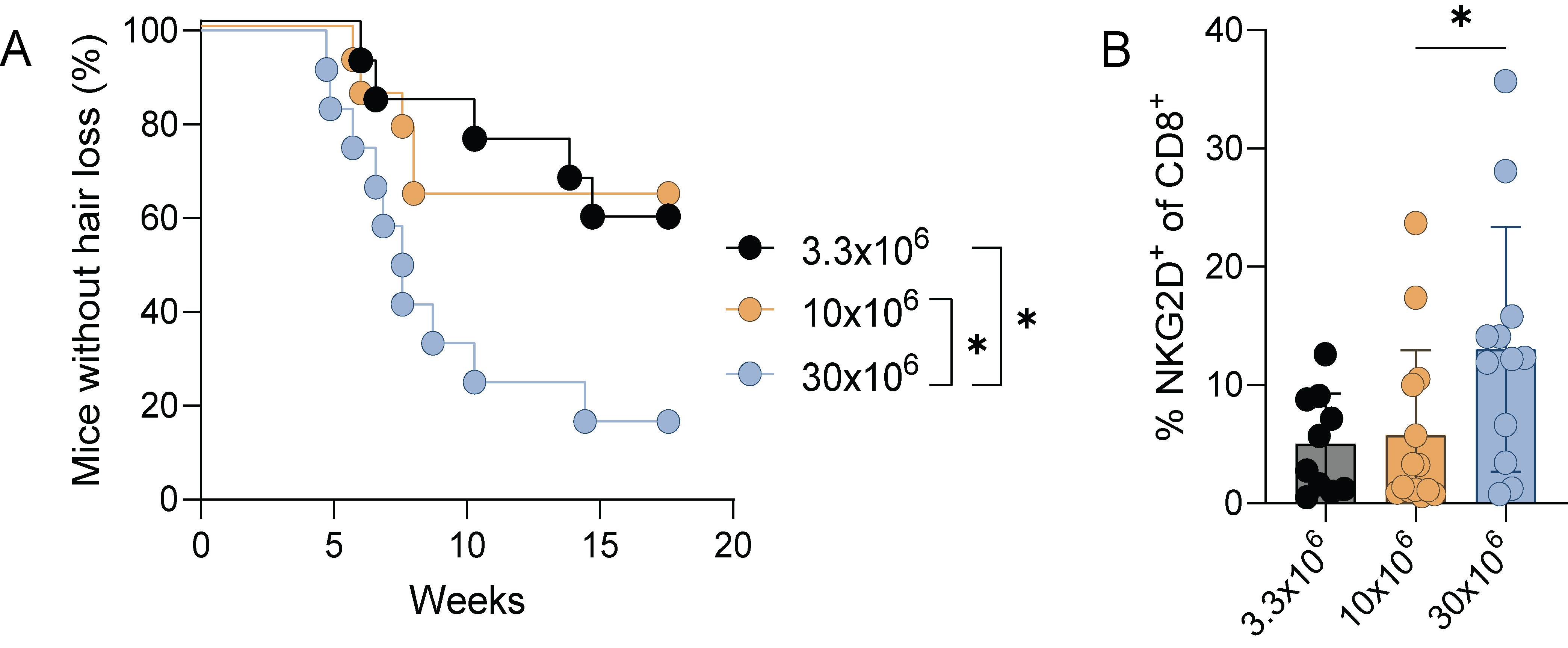

**Supplemental Figure 2. CD4 T cell mediated induction is dependent on the number of cells.** CD4 T cells were isolated from the SDLNs of AA donor mice and underwent *in vitro* expansion. Cells were intradermally injected into recipient mice at 3.3x10^6^ (n=12), 10x10^6^ (n=14), or 30x10^6^ (n=12) cells per mouse. **(A)** Kaplan-Meier curve for disease development. *p=0.0164 3.3x10^6^ vs 30x10^6^, *p=0.0165 10x10^6^ vs 30x10^6^, Log-rank test. **(B)** Frequency of NKG2D^+^CD8^+^ T cells in the SDLNs of recipient mice following disease induction (n=10 3.3x10^6^, n=14 10x10^6^, n=12 30x10^6^). *p=0.0455, Kruskal-Wallis test, *p=0.0451 30x10^6^ vs 10x10^6^, Dunn’s multiple comparison test. Two mice were excluded from the 3.3e^6^ group due to attrition immediately prior to analysis. Mice were observed two times weekly for the development of hair loss. Data representative of three combined experiments.

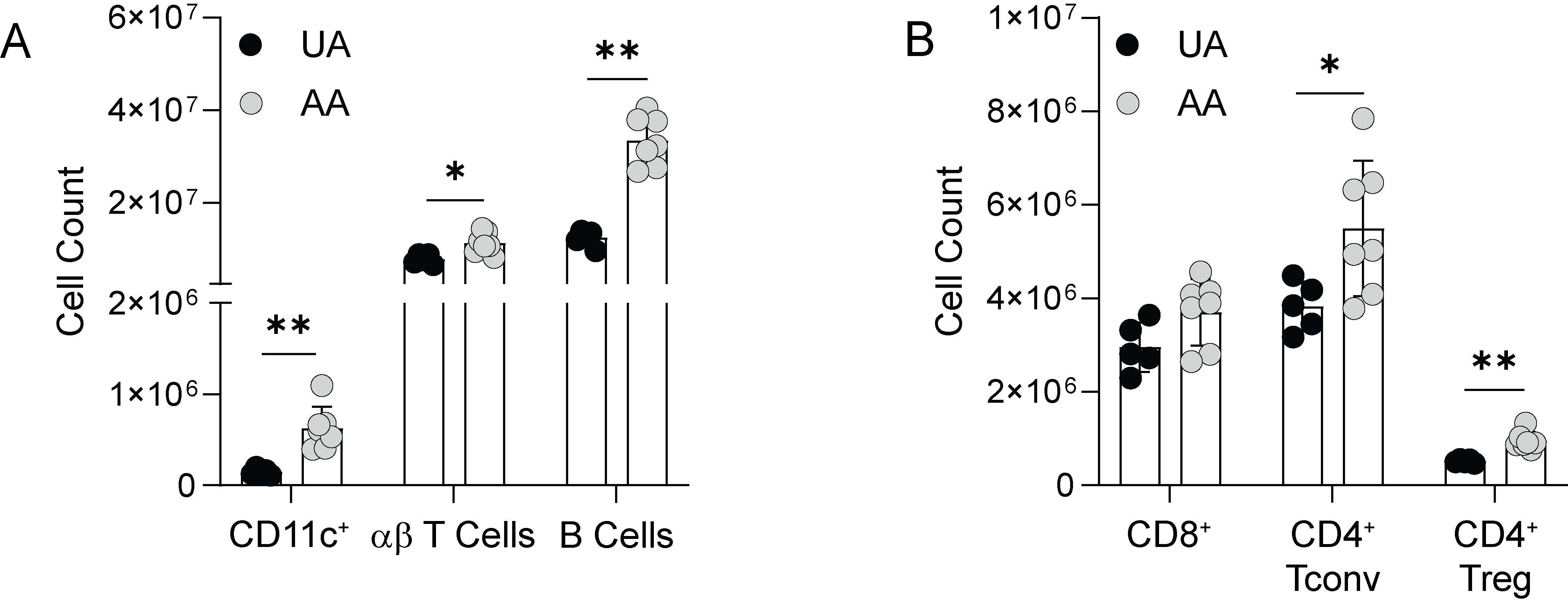

**Supplemental Figure 3. AA mice have an increased number of CD4 T cells in the spleen. (A)** Total number of CD11c^+^, αβ T cells (TCRβ^+^), and B cells (CD19^+^) in the spleens of UA and AA mice. **p=0.0025 CD11c^+^, *p=0.0101 αβ T cells, **p=0.0025 B cells, independent Mann-Whitney tests. **(B)** Total number of CD8^+^, CD4 Tconv (CD4^+^FoxP3^-^), and CD4 Treg (CD4^+^FoxP3^+^) T cells in the spleens of UA and AA mice. *p=0.0480 CD4 Tconv, **p=0.0025 Treg, independent Mann-Whitney Tests. Data is representative of two independent experiments.

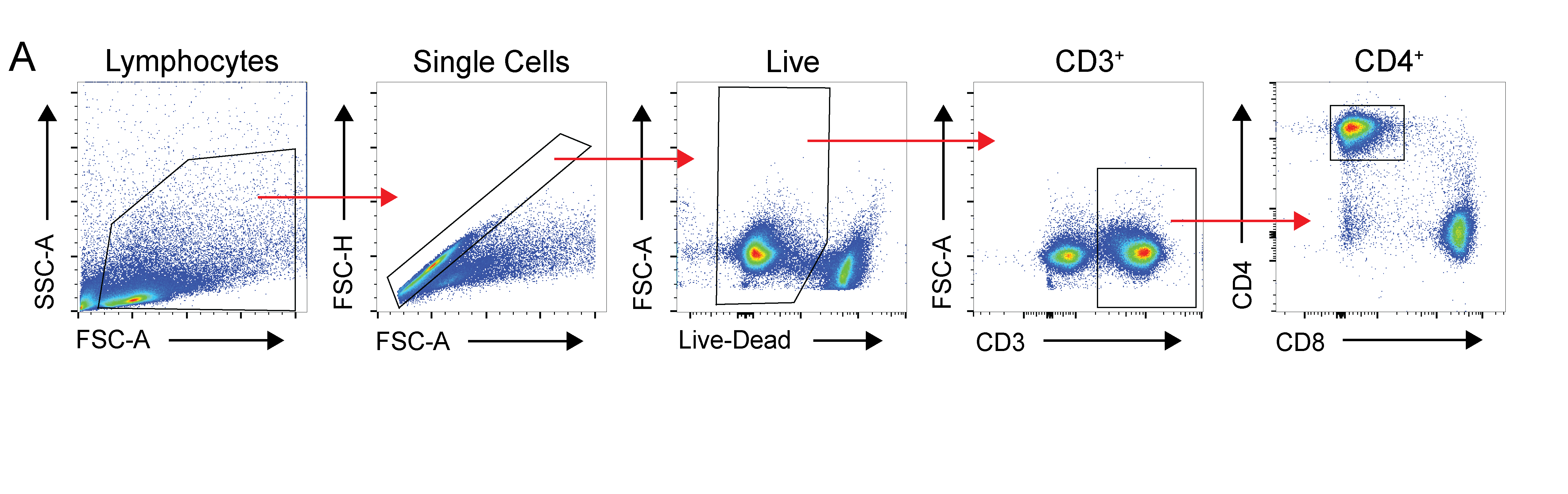

**Supplemental Figure 4. Gating strategy for conventional CD4 T cells for tSNE analysis.** (A) Representative gating strategy for tSNE analysis in Figure 4. Data is representative of at least two independent experiments.

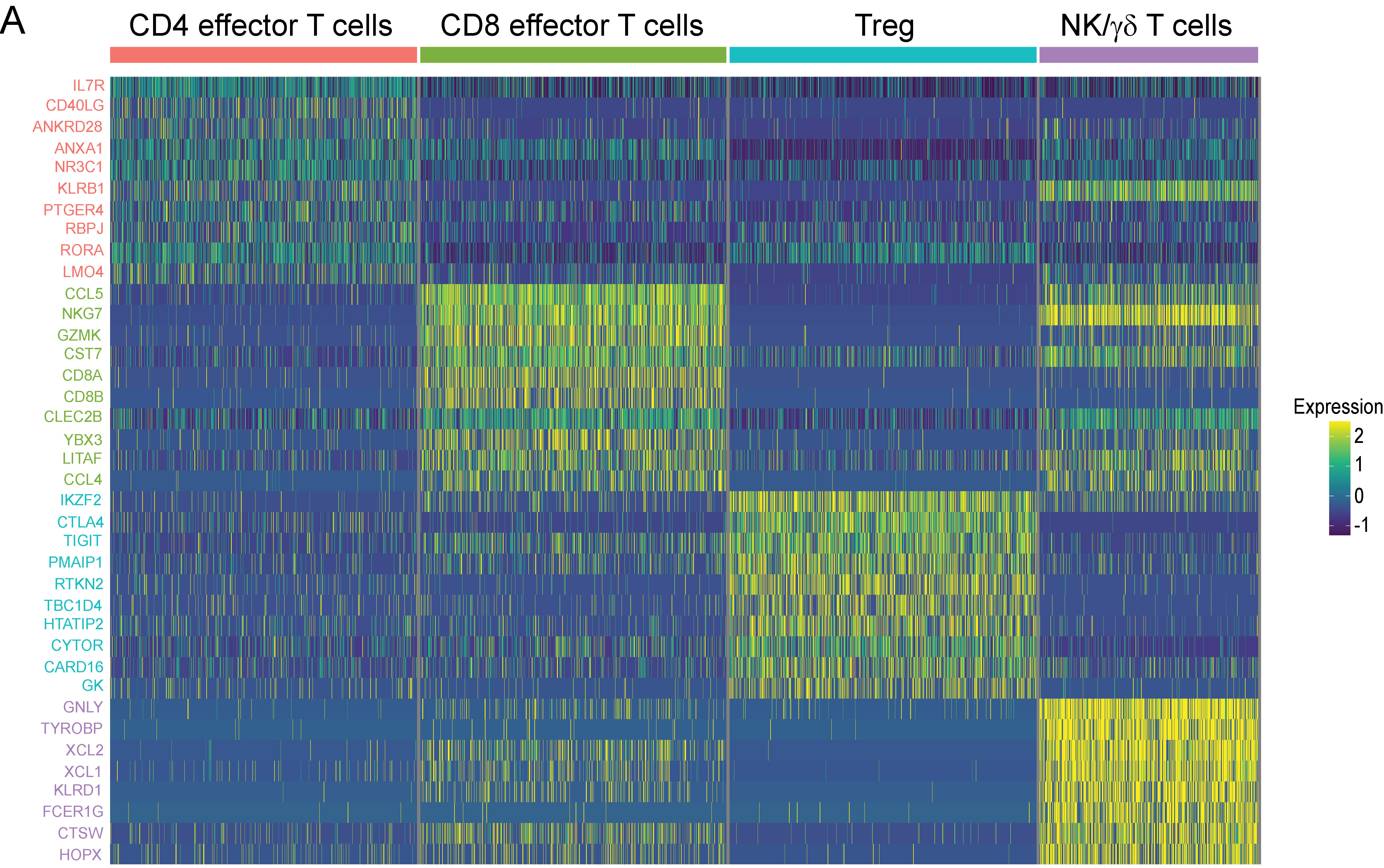

**Supplemental Figure 5. Human scRNA-seq T cell and NK cell identification. (A)** Heatmap of the top differentially expressed genes, ranked by percent difference, used to identify each of the 4 cell populations.
