## Supplementary material for "Th1 effector CD4 T cells rely on IFN-γ production to induce alopecia areata": Table 1 - Mouse Flow Antibodies

**Table 1: Mouse specific flow antibodies**

| **Marker** | **Fluorophore** | **Clone** | **Company** | **Catalog #** |
| --- | --- | --- | --- | --- |
| CD16/32 | - | 93 | Biolegend | 101302 |
| Viability | Live Dead Blue | - | Invitrogen | L34961 |
| Viability | Ghost Dye Violet 540 | - | Cytek Biosciences | 13-0879-T100 |
| Viability | Ghost Dye Red 780 | - | Cytek Biosciences | 13-0865-T100 |
| CD3ε | PE-Cy7 | 145-2C11 | Cytek Biosciences | 60-0031-U100 |
| CD3 | PE-Cy7 | 17A2 | Cytek Biosciences | 60-0032-U100 |
| CD3 | APC-Cy7 | 17A2 | Cytek Biosciences | 25-0032-U100 |
| CD4 | APC | GK1.5 | Cytek Biosciences | 20-0041-U100 |
| CD4 | APC | RM4-5 | Cytek Biosciences | 20-0042-U100 |
| CD4 | APC-Cy7 | RM4-5 | Biolegend | 100526 |
| CD4 | APC-Fire810 | GK1.5 | Biolegend | 100480 |
| CD8α | BUV615 | 53-6.7 | BD Biosciences | 613004 |
| CD8α | Spark UV 387 | 53-6.7 | Biolegend | 100798 |
| CD8α | PerCP-Cy5.5 | 53-6.7 | Cytek Biosciences | 65-0081-U100 |
| CD8α | PE-Cy5 | 53-6.7 | Cytek Biosciences | 55-0081-U100 |
| CD8α | FITC | 53-6.7 | Cytek Biosciences | 35-0091-U100 |
| TCRβ | FITC | H57-597 | Cytek Biosciences | 35-5961-U500 |
| TCRβ | APC | H57-597 | Cytek Biosciences | 20-5961-U100 |
| CD19 | red-Fluor710 | 1D3 | Cytek Biosciences | 80-0193-U025 |
| NKG2D | PE/Dazzle | CX5 | Biolegend | 130214 |
| NKG2D | PE | CX5 | Invitrogen | 12-5882-82 |
| CD11a | BUV661 | M17/4 | BD Biosciences | 741490 |
| CD44 | PerCP | IM7 | Biolegend | 103036 |
| CD44 | PE-Cy7 | IM7 | Cytek Biosciences | 60-0441-U100 |
| CD62L | BV605 | MEL-14 | Biolegend | 104438 |
| CXCR3 | BV421 | CXCR3-173 | BD Biosciences | 562937 |
| CD127 | PE-Cy5 | A7R34 | Biolegend | 135015 |
| CD28 | PE-Cy7 | 37.51 | Biolegend | 102125 |
| CD69 | FITC | H1.2F3 | Cytek Biosciences | 35-0691-U100 |
| CD25 | PE | PC61.5 | Cytek Biosciences | 50-0251-U100 |
| CD11c | PE | N418 | Biolegend | 117308 |
| FoxP3 | Pe-Cy5.5 | FJK-16S | Invitrogen | 35-5773-82 |
| I-A/I-E | BV711 | M5/114.15.2 | Biolegend | 107643 |
| IFN-γ | FITC | XMG1.2 | Cytek Biosciences | 35-7311-U100 |
| IFN-γ | APC | XMG1.2 | Cytek Biosciences | 20-7311-U100 |
| IFN-γ | BV711 | XMG1.2 | BD Biosciences | 564336 |
| IL-17A | PE | TC11-18H10.1 | Biolegend | 506903 |
