## Supplementary material for "Th1 effector CD4 T cells rely on IFN-γ production to induce alopecia areata": Table 2 - Human Flow Antibodies

**Table 2: Human specific flow antibodies**

| **Marker** | **Fluorophore** | **Clone** | **Company** | **Catalog #** |
| --- | --- | --- | --- | --- |
| Human TruStain FcX | - | - | Biolegend | 422302 |
| Viability | Live Dead Blue | - | Invitrogen | L34961 |
| Viability | Zombie UV | - | Biolegend | 423107 |
| CD3 | PE-Cy5 | UCHT1 | Biolegend | 300410 |
| CD4 | BV570 | OKT4 | Biolegend | 317445 |
| CD8α | BV711 | RPA-T8 | Biolegend | 301043 |
